# The crystal structure and localization of *Trypanosoma brucei* invariant surface glycoproteins suggest a more permissive VSG coat in the tsetse-transmitted metacyclic stage

**DOI:** 10.1101/477737

**Authors:** Aitor Casas-Sánchez, Samïrah Perally, Raghavendran Ramaswamy, Lee R. Haines, Clair Rose, Cristina Yunta, Marcela Aguilera-Flores, Michael J. Lehane, Igor C. Almeida, Martin J. Boulanger, Alvaro Acosta-Serrano

## Abstract

*Trypanosoma brucei* spp. develop into mammalian-infectious metacyclic trypomastigotes inside the tsetse salivary glands. Besides acquiring a variant surface glycoprotein (VSG) coat, nothing is known about expression of invariant surface antigens by the metacyclic stage. Proteomic analysis of saliva from *T. brucei*-infected flies revealed a novel family of hypothetical GPI-anchored surface proteins herein named Metacyclic Invariant Surface Proteins (MISP). MISP are encoded by five homolog genes and share ~80% protein identity. The crystal structure of MISP N-terminus at 1.82 Å resolution revealed a triple helical bundle that shares key features with other trypanosome surface proteins. However, molecular modelling combined with live fluorescent microscopy suggest that MISP N-termini are extended above the metacyclic VSG coat, exposing immunogenic epitopes. Collectively, we suggest that the metacyclic cell surface architecture appears more permissive than bloodstream forms in terms of expression of invariant GPI-anchored glycoproteins, which could be exploited for the development of novel vaccines against African trypanosomiases.

## Introduction

African trypanosomes are the causative agents of human African trypanosomiasis (HAT or sleeping sickness) and animal African trypanosomiasis (AAT or *Nagana*) in most sub-Saharan countries. Multiple trypanosome species, including the group of *Trypanosoma brucei* parasites (i.e. *T. b. gambiense*, *T. b. rhodesiense,* and *T. b. brucei*), are transmitted by several tsetse species (*Glossina* spp.). Although the number of cases of HAT have declined since 2010, due largely to a program of active screening and treatment of the human population together with an efficient vector control program, AAT continues to cause huge economic losses in the agricultural sector with >1 million cattle dying of AAT each year. In many cases, the disease may become fatal if untreated and current chemotherapies are generally toxic and inadequately administered^1^. Because the parasites undergo antigenic variation to effectively evade the host’s immune system, preventive vaccines are difficult to develop and none are yet available.

During its life cycle, *T. brucei* undergoes many developmental changes to adapt to the different environments within the mammalian host and the tsetse vector^2^. All *T. brucei* life stages have in common the expression of major surface glycosylphosphatidylinositol (GPI)-anchored glycoproteins known to be important for parasite survival. In bloodstream forms (BSFs), the variant surface glycoprotein (VSG) coat is responsible for antigenic variation^3–5^ and for the clearance of immunoglobulins against specific VSGs^6^. Furthermore, BSF VSGs form a macromolecular barrier that prevents binding of antibodies against conserved VSG epitopes and invariant (low copy number) surface proteins such as transporters and receptors^7–10^. When ‘stumpy’ BSF trypanosomes are ingested by the fly into the midgut (MG), they quickly (24 to 48 h) differentiate into the procyclic trypomastigote stage^2,11^. During differentiation, VSGs are no longer expressed and the parasites lose the VSG coat via the concerted action of major surface metalloproteases and the GPI-phospholipase C (GPI-PLC)^12,13^, which is a process that appears to modulate the tsetse’s immunity in favor of the parasite infection^14^. Concomitant with VSG disappearance, the parasites express a new coat of GPEET-procyclins, which is later replaced by a family of EP-procyclin isoforms once the infection is established in the MG^15,16,17^. It is well accepted that procyclic trypomastigotes colonize the tsetse ectoperitrophic space, from which they further migrate to the tsetse cardia or proventriculus (PV)^18^. Within the PV, mesocyclic trypomastigotes (MSCs) eventually differentiate into epimastigote forms (EMFs), migrate to the salivary glands (SGs), attach to the epithelial cell microvilli and switch on the expression of the surface-localized Brucei Alanine-Rich Proteins (BARP)^19^. The attached EMFs further differentiate into pre-metacyclic trypomastigotes (P-MCFs)^20,21^, during which time BARP expression is lost, a new coat of metacyclic VSG (mVSG) develops^22,23^, and cells detach from the SG epithelium. Finally, mature metacyclic trypomastigote forms (MCFs)^21,24^ are inoculated along with saliva into the skin of a vertebrate host in a subsequent feed. Unlike in BSFs, the surface composition of *T. brucei* MCFs is less well characterized primarily due to the lack of an *in vitro* culture system and to the low yield of MCF cells obtained from infected tsetse. Recently, however, trypanosomes overexpressing the RNA-binding protein 6 have been shown to develop into MCFs *in vitro*^25^, although the yields are low and the environment does not precisely mimic the tsetse SG.

It is well established that *T. brucei* BSFs have a large repertoire of *vsg* genes (>1000), but only one gene is expressed at a time^26,27^ from a single active expression site (BES), in which homologous recombination appears to exchange *vsg* genes and form mosaics^28^. In contrast, it is assumed that MCFs express a single mVSG (also called metacyclic variant antigen type; mVAT) from a much-reduced gene repertoire from the metacyclic expression sites (MES) that do not appear to recombine^22^. Based on antibody recognition, it is estimated that a population of tsetse-derived MCFs express different mVSGs (as many as 27) to ensure infection of hosts that may have been pre-exposed to any of these antigenic types^29,30^. Given that MCFs develop in the tsetse SGs and their persistence in the host after transmission is relatively short^24^, they seem not to be under pressure to undergo antigenic variation. Despite the differences in the expression mechanisms between VSGs and mVSGs, it is assumed that their function and structural organization is equivalent. For example, the BSF VSGs form a protective coat (composed of ~5 million homodimers/cell)^31^ that hides invariant surface proteins from the host immune system. Many of the known invariant proteins are predicted to be shorter in height than the VSG homodimers and to be attached to the plasma membrane via transmembrane domains^7,10^. Exceptionally, the haptoglobin-hemoglobin (HpHbR) and transferrin receptors are GPI-anchored and exclusively localize in the BSF flagellar pocket^8,9^. During a blood meal, the infected tsetse inoculates MCFs into the host along with saliva^24,32^ to inhibit the host hemostatic response to the cutaneous trauma^33,34^. However, during parasite development in the tsetse SGs, there is a drastic (~80%) transcriptional down regulation of genes encoding for salivary proteins, which induces a feeding phenotype that may amplify disease transmission^32,35,36^. Although the proteome of tsetse saliva from *T. brucei*-infected flies has been recently determined^34^, soluble parasite factors have not been characterized in detail and thus, the molecular mechanism by which these parasites manipulate the vector to promote transmission is largely unknown. Here we identify trypanosome proteins from infected tsete saliva using a high-resolution mass spectrometry-based proteomics approach. We found that *T. brucei*-infected saliva is particularly enriched with peptides from trypanosome surface proteins such as BARP, mVSG and a novel family of previously annotated hypothetical proteins we herein named Metacyclic Invariant Surface Proteins (MISP). We show that MISP are a small family of invariant surface glycoproteins that are primarily expressed in the metacyclic stage of *T. brucei*. Structural data reveal a triple helix bundle architecture for the MISP360 isoform that appears to be tethered to the outer membrane by an extended, unstructured C-terminal tail, projecting MISP above the mVSG coat and allowing antibody binding. Collectively, our findings provide a new understanding of the surface architecture composition of MCFs and open a new avenue for the potential generation of vaccines against different species of African trypanosomes.

## Results

### Saliva from *T. brucei-*infected tsetse is enriched with parasite GPI-anchored surface proteins

We initially investigated the protein composition of saliva from tsetse with a *T. b. brucei* infection in the SGs with the aim of identifying novel soluble factors that may be important for parasite transmission. We performed a high-resolution nano-liquid chromatography-tandem mass spectrometry (nLC-MS/MS) proteomic analysis of a pool of saliva from flies at 30 days post-infection (dpi), in three biological replicates (Supplementary Fig. 1). Having removed *Glossina* protein hits (Supplementary File 1), we detected in both naïve and trypanosome-infected saliva peptides from the bacterial symbiont *Sodalis glossinidius*^37^ (Supplementary File 2), whose presence was corroborated by immunoblotting using the anti-*Sodalis* Hsp60 1H1 monoclonal antibody (Supplementary Fig. 2). After manual curation of the MS/MS spectra, we identified 45 unique peptides derived from a total of 27 *T. b. brucei* proteins in infected saliva datasets, with high confidence identification (>95% at protein level) (Supplementary File 3). Of the identified proteins, 17 (62.9%) are predicted to be intracellular (Supplementary Table 1) mainly linked to functions such as protein folding (14.8%), cytoskeleton (11.1%), and metabolic processes (7.4%). The remaining 10 proteins (37.1%), are predicted to be GPI-anchored surface proteins from three major families (Table 1). Eight putative BARP isoforms were identified from four unique peptides, and five different VSGs were identified from seven peptides (Table 1, Supplementary File 4). Notably, we identified a single peptide (SVAEDNSAASTAR) with 100% confidence (supported by two spectra, Supplementary File 4), which belongs to the small family of hypothetical GPI-anchored MISP (Table 1). This peptide is fully conserved across all *T. brucei* MISP isoforms (Fig. 1b, Supplementary File 4).

**Table 1.** Summary of major GPI-anchored proteins from *T. b. brucei* identified in infected tsetse saliva by nLC-MS/MS. All identified proteins and peptides have >95% confidence and were manually curated. Protein accession code (Protein ID), protein description (Annotation), number of identified unique peptides (Peptides), and number of spectra (Spectra) are indicated.

| Protein ID <sup>(a)</sup> | Annotation | Peptides | Spectra <sup>(b)</sup> |
| --- | --- | --- | --- |
| Tb927.9.15520 <sup>(1)</sup> | Brucei Alanine-Rich Protein (BARP) | 1 | 3 |
| Tb927.9.15530 | Brucei Alanine-Rich Protein (BARP) | 4 | 7 |
| Tb927.9.15570 | Brucei Alanine-Rich Protein (BARP) | 1 | 2 |
| Tb927.9.15630 | Brucei Alanine-Rich Protein (BARP) | 1 | 1 |
| VSM2_TRYBB | Variant Surface Glycoprotein (VSG) 221 | 3 | 4 |
| O76421_TRYBR | Metacyclic Variant Antigen Type (mVAT) 4 | 1 | 1 |
| A0A1J0RB71_9TRYP | Variant Surface Glycoprotein (VSG) 1125.4959 | 1 | 1 |
| A0A1J0R978_9TRYP | Variant Surface Glycoprotein (VSG) 1125.3088 | 1 | 1 |
| A0A1J0RAQ3_9TRYP <sup>(2)</sup> | Variant Surface Glycoprotein (VSG) 1125.408 | 1 | 1 |
| Tb927.7.360 <sup>(3)</sup> | Metacyclic Invariant Surface Protein (MISP) | 1 | 2 |

<sup>(a)</sup> Accession code for *T. b. brucei* in TriTrypDB (MISP and BARP) or UniProt (VSG). <sup>(b)</sup> See peptides and spectra in Supplementary File 4. <sup>(1)</sup> Representative ID. Peptide conserved in Tb927.9.15520, Tb927.9.15530, Tb927.9.15550, Tb927.9.15590, Tb927.9.15600, Tb927.9.15620 and Tb927.9.15660. <sup>(2)</sup> Representative ID. Peptide also conserved in *T. b. brucei* VSGs 646, 769, 3613, 1125.408, 1125.474, 1125.1142, 1125.4207 and 1125.4707. <sup>(3)</sup> Representative ID. Peptide conserved across all 5 *T. brucei* MISP isoforms.

**Fig. 1.**
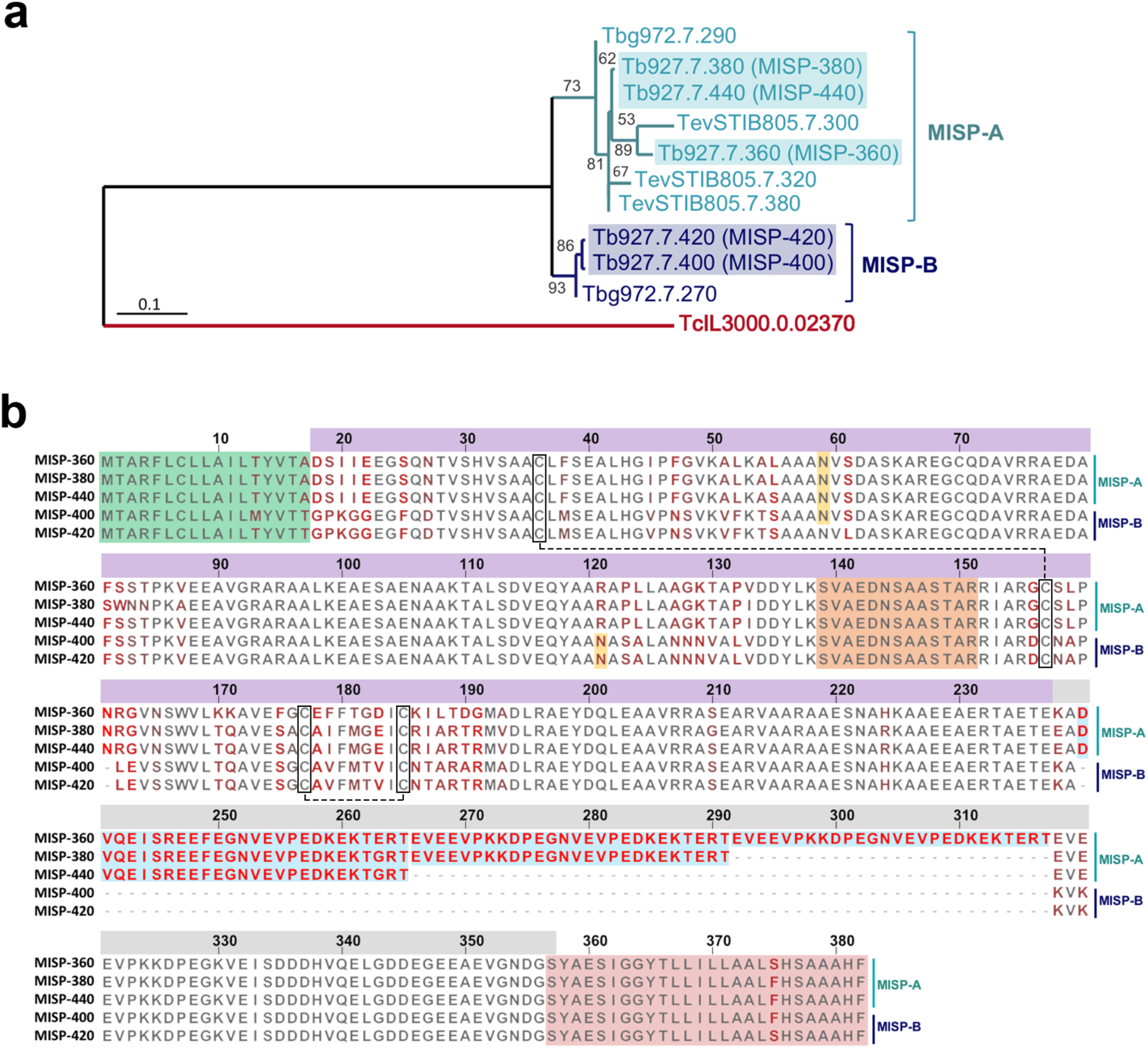
MISP sequence analyses. **a**, Unrooted phylogenetic tree generated from a multiple sequence alignment of all MISP isoforms. Note the demarcation of two sub-families MISP-A (green) and MISP-B (blue) and the least-conserved *T. congolense* MISP (red). The *T. b. brucei* MISP are highlighted. Tree created using PhyML 3.1 and the WAG substitution model of maximum likelihood; bootstrap of 500; numbers show branch support values (%); scale bar: 0.1 substitutions/site. **b**, Multiple protein alignment of the *T. b. brucei* MISP isoforms (MISP360, MISP380, MISP440, MISP400 and MISP420). Residues colored from grey (full conservation) to bright red (highest divergence). Protein domains highlighted: N-signal peptide (green), predicted *N-*glycosylation sites (yellow asparagine), disulphide bonds (cysteines in black boxes), C-terminus motifs of 26 residues (blue) and GPI-anchor attachment signal peptide (red). The conserved peptide ‘SVAEDNSAASTAR’ identified by nLC-MS/MS is highlighted in orange. The top bar numbers indicate residue position starting from Met^1^. Top bar colors indicate the main protein domains N-terminus (purple) and C-terminus (grey). In addition to the presence of characteristic sequences defining MISP-A and MISP-B families (see text and Supplementary Fig. 3a), other N-terminal residues further define each protein sub-family (e.g. Ser^25^, Asn^27^, Phe^38^, Arg^121^, and Pro^123^-Leu^124^ (for MISP-A), and Phe^25^, Asp^27^, Met^38^, Asn^121^, and Ser^123^-Ala^124^ (for MISP-B)).

### The Metacyclic Invariant Surface Proteins (MISP)

MISP proteins are part of the large Fam50 family of trypanosome surface proteins^38^. *T. b. brucei* contains five *misp* homolog genes (*Tb927.7.360, Tb927.7.380, Tb927.7.400, Tb927.7.420, Tb927.7.440* in strain TREU927), which encode for proteins that are unique in sequence and have no predicted function. All five *T. b. brucei* homologs are localised to chromosome 7 (Supplementary Fig. 3) and are separated by unrelated genes (*Tb927.7.370, Tb927.7.390, Tb927.7.410* and *Tb927.7.430*; all predicted to be homologs encoding for the Golgi complex component 7). The same *misp* homologs are found in the human pathogenic subspecies *T. b. rhodesiense* and *T. b. gambiense* (*Tbg972.7.270* and *Tbg972.7.290*). Furthermore, slightly divergent versions of *misp* genes can also be found in the animal trypanosome species *T. evansi* (*TevSTIB805.7.300, TevSTIB805.7.320* and *TevSTIB805.7.380*) and *T. congolense* (*TcIL3000.0.02370*). No homologs were found in the *T. vivax* genome. Using a phylogenetic analysis (PhyML 3.1)^39^, we determined that MISP are comprised of two sub-families (MISP-A and MISP-B) that originate from a common ancestor (Fig. 1a). Members of the MISP-A sub-family (*T. b. brucei* Tb927.7.360, Tb927.7.380, Tb927.7.440, *T. b. gambiense* Tbg972.7.290 and *T. evansi* TevSTIB805.7.300, TevSTIB805.7.320 and TevSTIB805.7.380) contain the N-terminal sequences Asp^18^-Ser-Ile-Ile-Glu^22^ and Ala^127^-Gly-Lys-Thr^130^ in addition to one to three repeats at the C-terminus (see below). In contrast, MISP-B isoforms (*T. b. brucei* Tb927.7.400 and Tb927.7.420, and *T. b. gambiense* Tbg972.7.270) harbor a conserved Gly^18^-Pro-Lys-Gly-Gly^22^ and Asn^127^-Asn-Asn-Val^130^ at the N-terminus and no C-terminal repeats. *T. congolense* TcIL3000.0.02370 cannot be included within either group as it does not share the common ancestor and it only has 34.2% identity with other MISP isoforms (Supplementary Fig. 3b). A comparison of the five *T. b. brucei misp* homologs revealed an overall amino acid sequence identity of ~79% (Fig. 1b).

In addition, the amino acid sequence alignment showed that the N-termini domains (Met^1^– Glu^237^) are more conserved (88.7% identity) than the C-termini (Lys^238^-Ser^357^), where the main divergence is found (49.0% identity). However, this divergence is not random but primarily localized to charged 26 residue repeats (EVEEVPKKDPEGNVEVPEDKEKTERT or DVQEISREEFEGNVEVPEDKEKTERT). Alignment of all MISP isoforms across species (excluding TcIL3000.0.02370) shows an overall identity of 79.1% (Supplementary Fig. 3a). Using the SignalP 4.1 server^40^, we identified signal peptides in all *T. b. brucei* MISP isoforms (Met^1^–Ala/Thr^17^), although, MISP-B Tb927.7.400 may also contain an alternative cleavage site at Gly^24^ (Supplementary Table 2). Notably, MISP from other trypanosome species have equivalent signal peptide predictions (Supplementary Table 2). While no transmembrane domains were identified for MISP (Supplementary Table 2), all *T. brucei* isoforms have a strongly predicted (99.9% specificity) GPI site^41^, with a putative omega site at Ser (underlined) within the C-terminal GPI addition sequence <u>S</u>YAESIGGYTLLILLAALS/FHSAAAHF (Fig. 1b, Supplementary Table 2). Furthermore, the NetNGlyc 1.0 server predicts that all MIS Phomologs contain a potential *N-*glycosylation site at Asn^58/59^, except for Tb927.7.420 (with no site) and TcIL3000.0.02370 (seven sites) (Supplementary Table 2).

### Developmental regulation of *misp* transcripts

To better understand the expression pattern of *misp* during parasite development in the tsetse, we used semi-quantitative and real time RT-PCR, alongside detection of *barp* as a marker for SG EMFs^19^. Total RNA samples were isolated from all *T. b. brucei* life stages obtained from both *in vitro* cultures (i.e., BSFs and procyclic cultured forms (PCFs)) and tsetse-derived parasites at 30 dpi. (i.e., MG procyclics, PV parasites (95% MSCs, 5% EMFs), whole infected SGs, and isolated MCFs). Samples from naïve flies (30 days old) were used as negative control. Due to the high sequence identity among the five *misp* homolog genes, assessing their individual expression by real-time RT-PCR was not possible. We therefore used the more permissive semi-quantitative RT-PCR method and exploited the differential lengths of the C-terminal repeats, which are unique for four of the five homologs and show differences of 78 bp (Supplementary Fig. 5a). While control samples from naïve flies were all negative for *misp*, there is a significant up regulation of *misp* in SG stages compared to MG (11.8-fold) and BSF (25.1-fold) stages, with p-values <0.001 (Fig. 2a). Although isolated MCFs also presented similar expression levels compared to SGs (1.5-fold less, p=0.003), we could not rule out that ~5% contaminating EMFs in saliva also contributed to these high expression levels. PV trypanosomes, despite having a 2.1-fold (p<0.001) lower expression compared to SG stages, showed significantly higher expression than MG procyclics, BSFs, and PCFs (>5.5-fold, p=0.002), which together express negligible levels of *misp* with no significant differences between them. Although there seems to be an overall preferential expression for homologs with short C-termini (*misp400/420*), the longer homologs *misp380* and *misp360* were only expressed in SG and MCF stages. To further validate the robustness of our detection method, we analyzed the expression of *misp* in a mutant PCF cell line overexpressing an ectopic copy of *misp360* under a doxycycline-inducible system. While uninduced (Dox-) control cells showed *misp* expression levels similar to wild type PCF, induced (Dox+) cells showed a 2.8-fold increase in *misp*360 only (Supplementary Fig. 5a). On the other hand, the expression levels of *barp* (all 14 homologs share >60% identity at protein level, Supplementary Fig. 4) were also determined using the same samples for comparison (Fig. 2b). *Barp* transcripts were found to be up regulated in PV parasites compared with MG procyclics (10.6-fold), PCFs (24-fold), and BSFs (26.6-fold), with p-values <0.01. This high expression was maintained in SG stages, including MCFs. To validate these results, a similar expression analysis was performed using real time RT-PCR targeting universal sequences within the *misp* and *barp* coding sequences (Supplementary Fig. 5b). The same trend of high expression was observed for both *misp* (40-fold) and *barp* (95.6-fold) in SG compared to MG isolated parasites, although with a greater fold change.

**Fig. 2.**
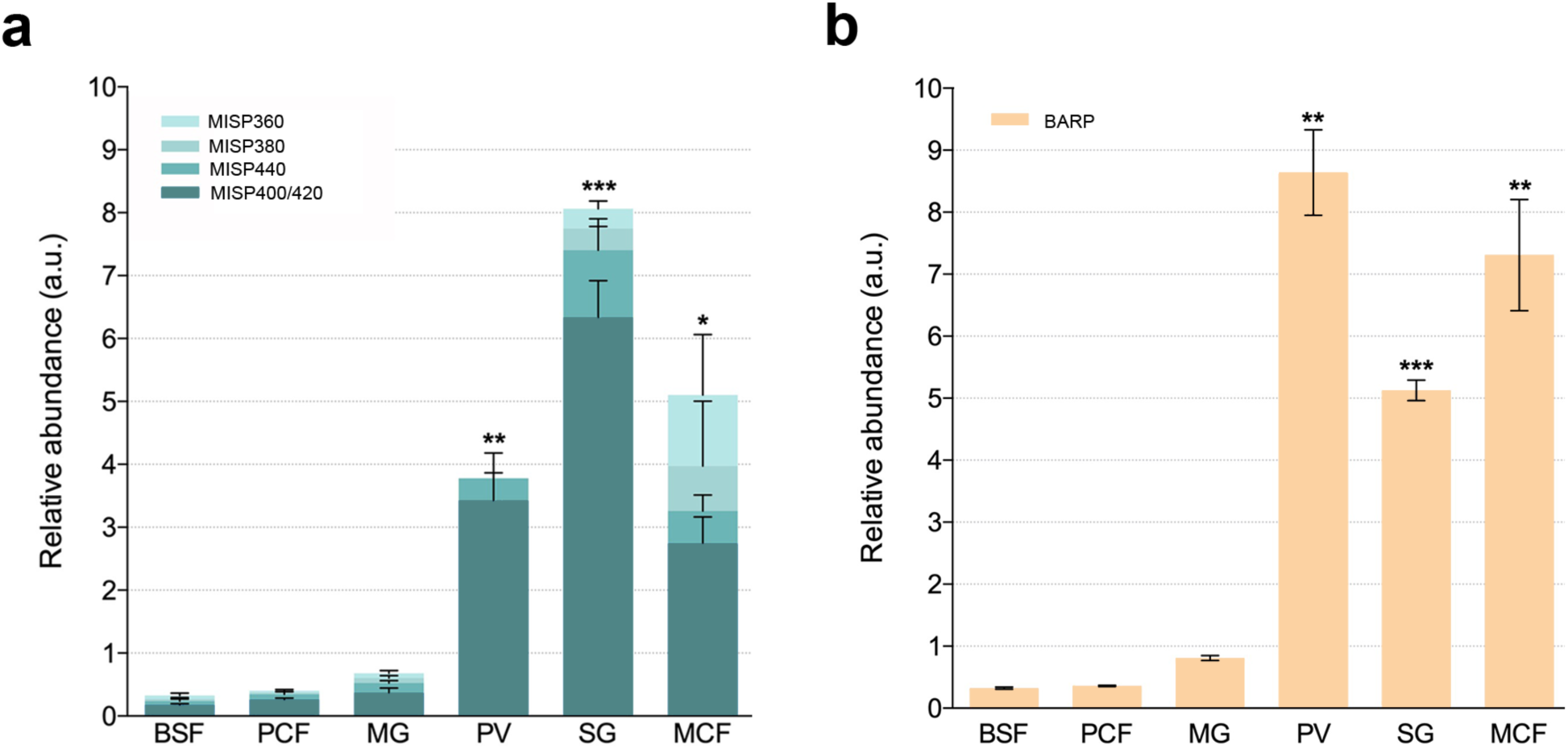
Expression levels of *misp* and *barp* throughout the *T. b. brucei* life cycle. **a**, Relative mRNA expression levels (arbitrary units, normalised to the expression of *tert*) of the *T. b. brucei misp* homologs, detected by semi-quantitative RT-PCR targeting the C-termini motifs in *T. b. brucei* AnTat 1.1 90:13 cultured bloodstream forms (BSF), procyclic cultured forms (PCF), midgut procyclics (MG), proventricular forms (PV), salivary gland forms (SG) and isolated metacyclic forms (MCF). **b**, Relative mRNA expression levels of *T. b. brucei barp*, targeting a universal region found in all homologs. Error bars represent standard deviation, asterisks show statistical significance (* for p-value <0.05, ** <0.01, *** <0.001).

### MISP are homogenously expressed on the surface of *T. brucei* salivary gland stages

Having identified SG stages as the major expressors of *misp* and *barp*, we next determined their localizations by immunofluorescence using polyclonal antibodies, which recognize conserved epitopes across the MISP or BARP isoforms. As predicted, recognition only occurred in infected SGs (Fig. 3) and neither BSFs, PCFs, MG procyclics nor PV parasites were recognized by either antibody (all PV forms express exclusively EP-procyclin as major surface GPI-anchored protein, see Supplementary Fig. 6). Both polyclonal anti-MISP and anti-BARP antibodies recognized all parasite stages in infected SGs, staining both clustered cells (EMFs and P-MCFs) that appeared attached to the SG epithelium and free-swimming MCFs.

**Fig. 3.**
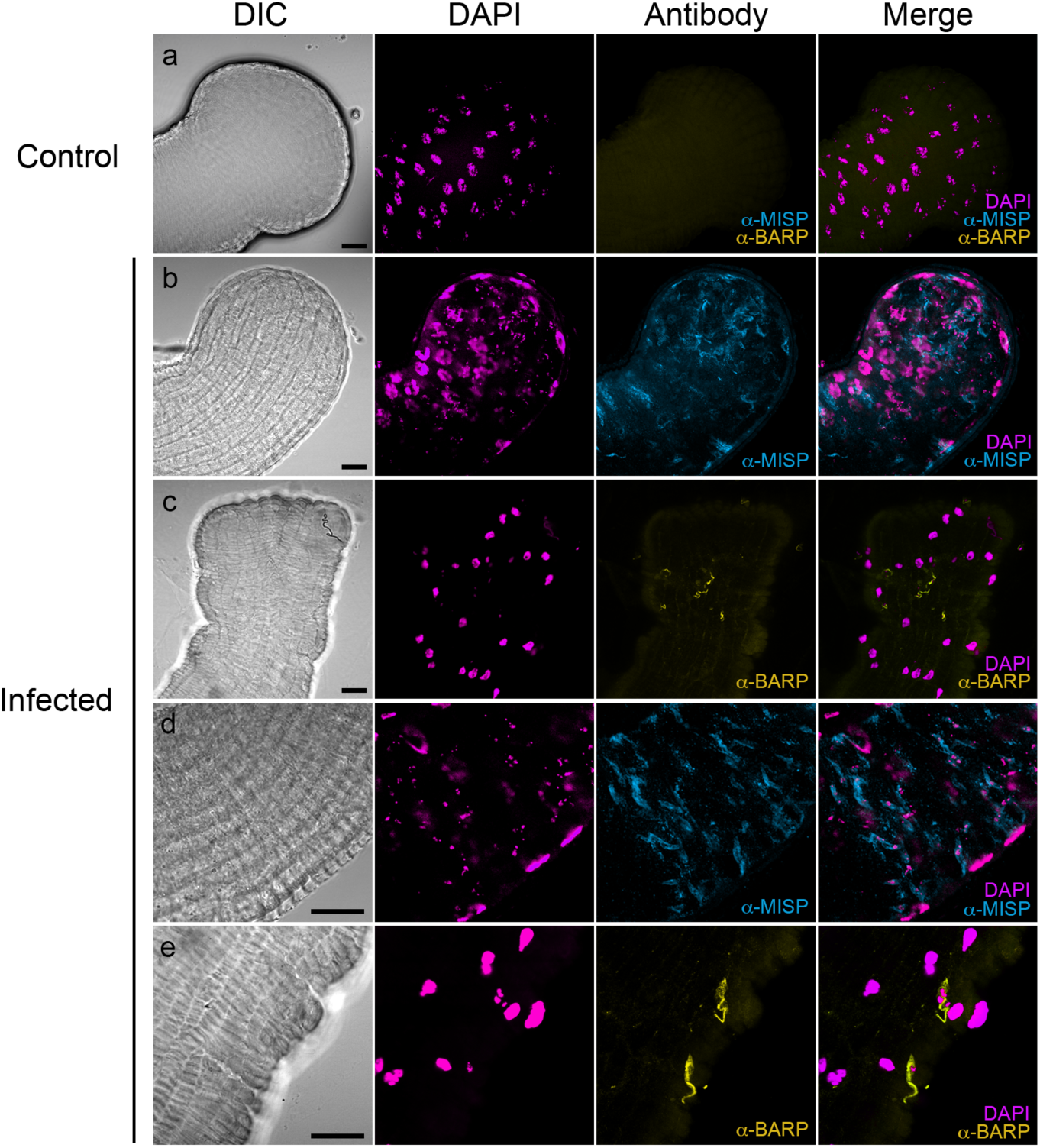
*T. b. brucei* salivary gland life stages express MISP and BARP during an infection in the tsetse salivary glands. Immunostaining of MISP and BARP proteins in trypanosomes infecting a tsetse salivary gland. Representative images of uninfected (a) and *T. b. brucei*-infected salivary glands (b-e). Differential interference contrast (DIC), DAPI (blue), anti-MISP (red) and anti-BARP (green). Scale bars: 20 μm.

To determine the precise cellular localization of these proteins, we probed individual parasites isolated from infected SGs. All EMFs, P-MCFs and MCFs reacted with anti-MISP and anti-BARP antibodies (Fig. 4a, Supplementary Fig. 7). The relative mean fluorescence intensity of MISP was higher in MCFs compared to P-MCFs (1.5-fold) and EMF (2.3-fold) (p-values <0.01) (Fig. 4b). This suggests that MISP expression starts in EMFs and progresses as parasites develop within the SGs, with the maximum expression found in mature MCFs. MISP appeared to localize evenly across the cell surface in non-permeabilized fixed cells (Fig. 4a). To confirm surface exposure of MISP epitopes, we performed immunostaining on live MCFs (Supplementary Fig. 8). All live cells stained uniformly when incubated with anti-MISP antibody, which confirms that at least part of MISP N-terminal epitopes are accessible to antibodies. Interestingly, when an eGFP and HA epitope-tagged MISP360 was ectopically expressed in ^HA-eGFP^MISP360 Dox+ PCFs (Supplementary Fig. 9a), the protein was exclusively detected in the flagellum as suggested by its co-localization with the flagellar marker, paraflagellar rod (PFR) (Supplementary Fig. 9b). However, when the same MISP360 overexpressing cell line progressed through the tsetse and colonized the SGs, the ectopic protein re-localized evenly on the MCF surface (Supplementary Fig. 9b), resembling that of the wild type MCFs. We then compared MISP expression and localization in relation to BARP. Using the same method, BARP was detected on the surface of non-permeabilized EMFs, P-MCFs and MCFs (Fig. 4c). However, the relative mean fluorescence intensities in EMFs were significantly higher compared to P-MCFs (1.3-fold) and MCFs (3.3-fold) (Fig. 4d), with p-values of 0.04 and <0.001, respectively. BARP proteins localize across the entire cell surface in EMFs and P-MCFs. However, in MCFs, surface expression of BARP was only detected in 10.4% of cells (n=481), while most of it (89.6%) appear to be flagellar (Fig. 4c, Supplementary Fig. 6). To confirm the identity of SG trypanosomes, cells were probed with anti-CRD polyclonal antibodies, which recognize the cross-reacting determinant (CRD) epitope formed after cleavage by the GPI-PLC^21,42^. While EMFs and P-MCFs did not react with the antibody, mature MCFs showed surface staining as an indirect detection of mVSGs (Supplementary Fig. 10).

**Fig. 4.**
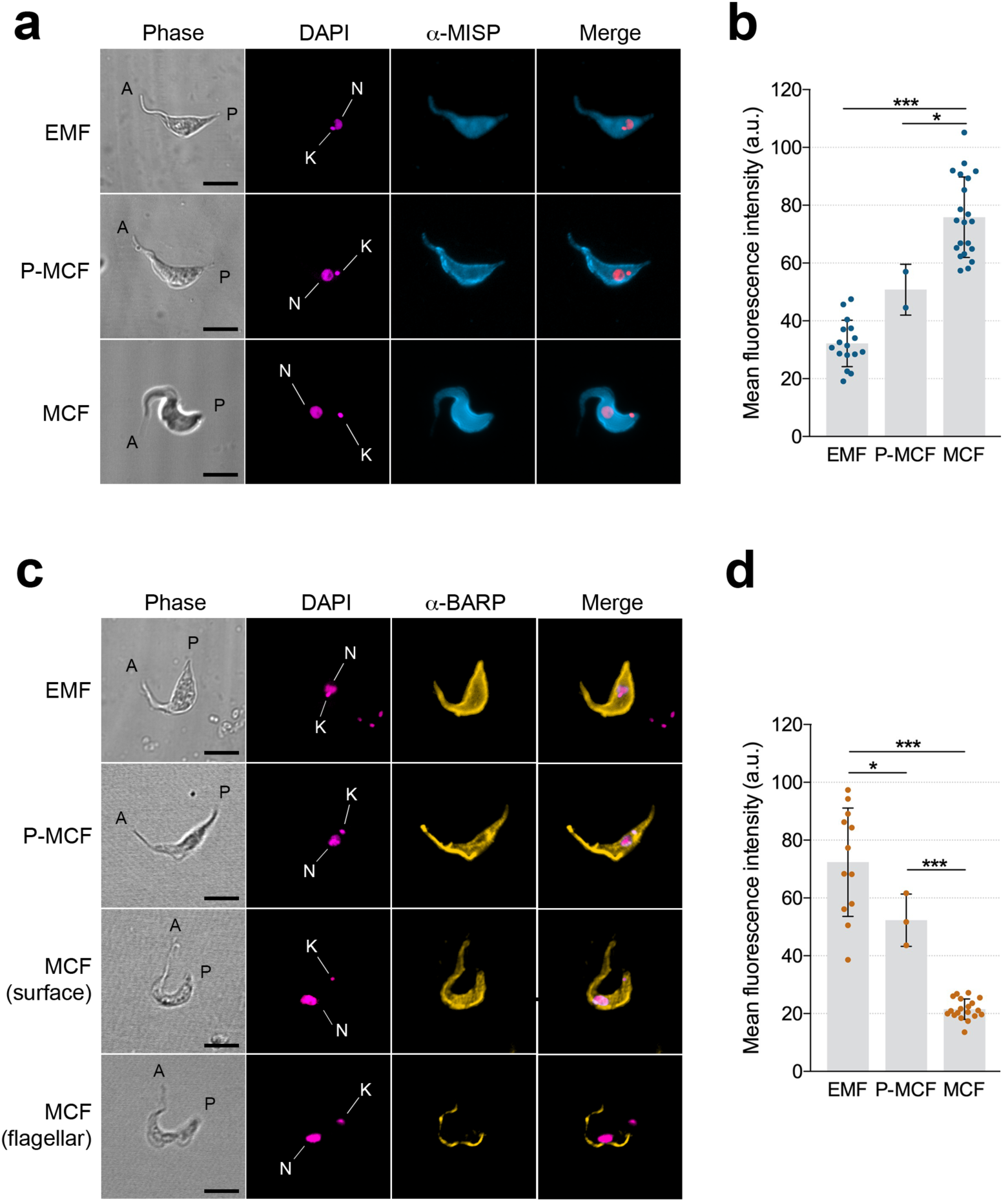
Expression and localization of MISP and BARP in *T. b. brucei* salivary gland stages. **a**, Localisation of MISP. Immunostaining of non-permeabilised parasites extracted from infected salivary glands: epimastigote (EMF), pre-metacyclic form (P-MCF) and metacyclic form (MCF). Phase, antibody anti-MISP (red), and DAPI (blue). Kinetoplasts (K) and nuclei (N) are highlighted in the blue channel. **b**, Mean fluorescence intensity (arbitrary units) of anti-MISP stained cells. **c**, Localisation of BARP. Immunostaining of non-permeabilised salivary gland parasites. Phase, antibody anti-BARP (green) and DAPI (blue). Representative MCFs for BARP surface and flagellar localisation. **d**, Mean fluorescence intensity (arbitrary units) of anti-BARP stained cells. Scale bars: 5 μm. Error bars show standard deviation, asterisks represent significance (* for p-value <0.05; *** for p-value <0.001). Neither anti-BARP nor anti-MISP recognized BSFs, PCFs, MG and PV forms (not shown).

### The N-terminal region of MISP adopts a triple helix bundle structure

To complement the expression and cellular localisation data, we next determined the crystal structure (1.82 Å resolution) of the conserved N-terminal ectodomain (Gly^24^–Ala^234^) of *Tb*MISP360 (Tb427.07.360) (Fig. 5a). It should be noted that Tb427.07.360 is an isoform of Tb927.7.360 in the strain Lister 427 of *T. b. brucei* and shares 98.7% sequence identity across the N-terminal domain (residues Met^1^ to Glu^237^). Crystals of purified, monomeric *Tb*MISP360 (Supplementary Fig. 11) were obtained using the hanging drop method and grew in space group P2_1_2_1_2_1_ with a single molecule in the asymmetric unit. The structure of *Tb*MISP360 was determined by molecular replacement using a truncated form of *T. congolense* GARP (GARP, PDB entry: 2Y44) as the search model. The identification of GARP as a suitable model was based on secondary structure predictions as the amino acid sequence identity of the N-terminal ectodomain is only 15%. The overall structure of *Tb*MISP360 was well defined with only two residues from the N-terminus remaining un-modelled (Gly^24^ and Ser^25^). The core of the N-terminal ectodomain adopts an elongated structure measuring approximately 83 Å in height and spanning approximately 25 Å in width (Fig. 5b, left). The ectodomain is well ordered with low *B-*factors throughout the protein (Fig. 5b, middle). Like GARP, *Tb*MISP360 adopts an overall triple helical bundle structure composed of a core of extended twisted helices capped by a smaller helical bundle at the N-terminal end. The helical bundle that dominates the structure consists principally of three helices (Fig. 5b, right): helix I is comprised of residues Val^29^-Ser^83^, helix II of Glu^88^-Thr^130^ and helix VI of Phe^182^-Ala^233^. The three helices adopt a bend of approximately 30° at Gly^44^ (helix I), Ala^127^ (helix II) and Leu^195^ (helix VI) and collectively, give rise to the helical bundle cap. In addition to the ends of the 3 major helices, this bundle includes 3 shorter helices: helix III (Asp^134^-Glu^142^), which is connected by a 5-residue loop to helix IV (Ser^148^-Gly^156^), and helix V (Ser^166^-Phe^179^) connected to helix IV by a 9-residue loop. Helices IV and V lie at either side of helix I and form the broadest face of the helical bundle cap.

The lack of significant sequence identity between *Tb*MISP360 and any protein with known function led us to perform a DALI search^43^ to identify structural homologs. GARP was identified as the top hit with a Z-score of 21.4. A least squares superposition between the two structures resulted in a root-mean-square deviation of 1.7 Å over 183 Cα atoms (Fig. 5c, left). The N-terminal domain of GARP has two long helices and a third, smaller helical strand that runs back along the helical bundle. In *Tb*MISP360, the helices adopt a similar pattern with the third helix having an additional 11 amino acids when compared to GARP. Moreover, the head structure of *Tb*MISP360 is anchored by two disulphide bonds; one between Cys^36^ and Cys^157^, and the other between Cys^177^ and Cys^185^ (Fig. 1b), whereas GARP incorporates only one disulphide bond between Cys^48^ and Cys^161^. The DALI search also revealed structural homology to the previously characterized HpHbR from *T. brucei* (PDB: 4X0J) and *T. congolense* (PDB: 4E40)^44,45^, and to a *T. brucei* VSG monomer (PDB: 1VSG) with Z-scores of 18.1, 17.7 and 6.5, respectively. All these proteins exhibit a complementary core of twisted three helical bundles (Supplementary Fig. 11). Structural comparison, however, indicates a closer architectural similarity with HpHbR compared to VSG. This is due, in part, to the breakdown of the third helix into loops and extensions enabling substantial conformational diversity in VSG^46^. Moreover, *Tb*MISP360 is monomeric in contrast to dimeric VSGs^46^.

Despite the general structural similarity with GARP, *Tb/Tc*HpHbR and the VSG monomer, the overall low sequence identity and the lack of key conserved residues suggest a different biological role for MISP.

**Fig. 5.**
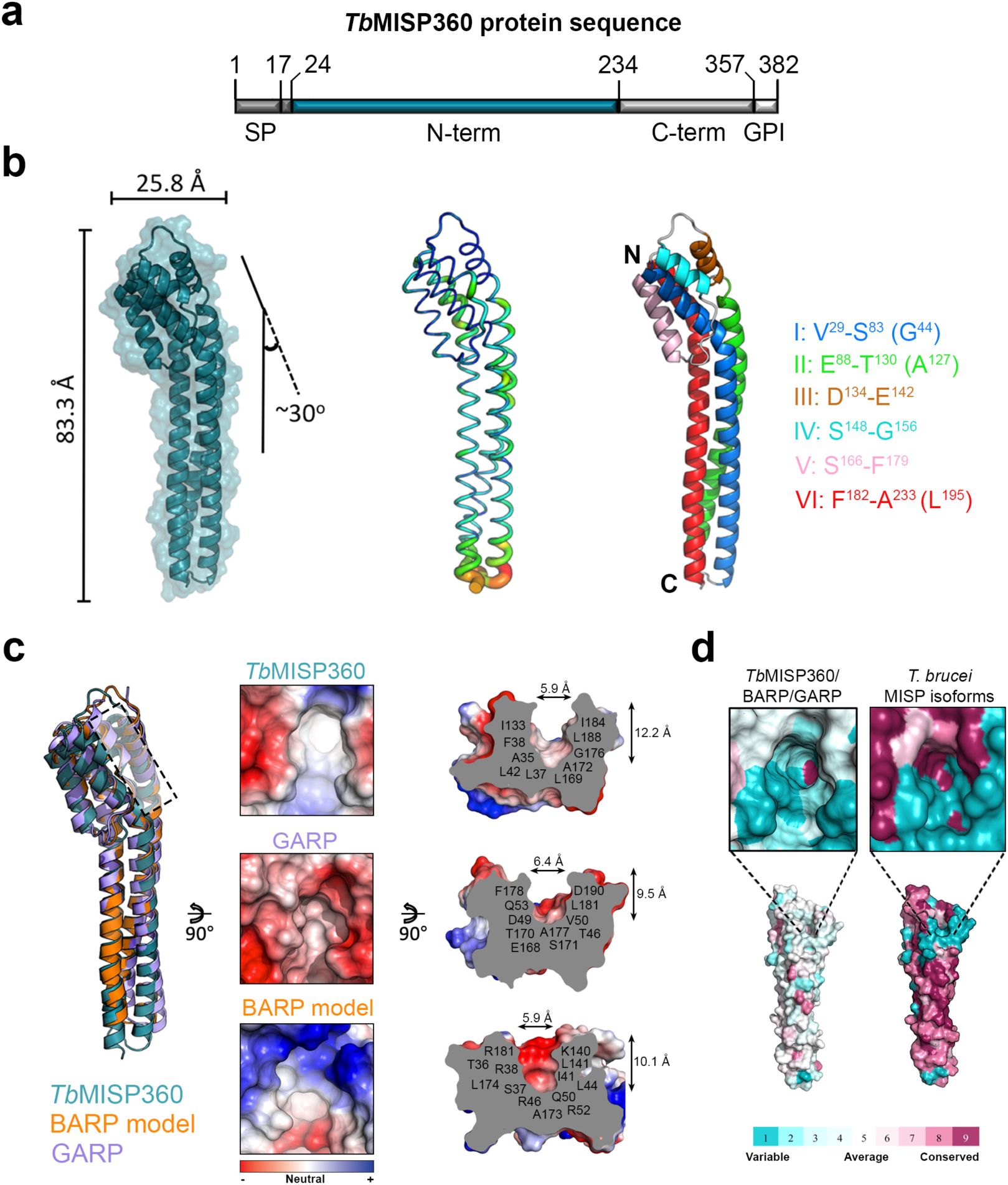
Crystal structure of the MISP N-terminal ectodomain. **a**, Construct encoding the *Tb*MISP360 N-terminus that was recombinantly expressed in *E. coli* and crystallised (Asn^24^-Glu^235^). **b**, *Tb*MISP360 N-terminus crystal structure with surface representation in purple. The structure was found to be 83.3 Å tall, 25.5 Å in width with a 30° helix bend at the top (left); *B-*factor putty model with ordered regions displayed in blue and thin tubes and flexible regions in red and thicker tubes (middle); secondary structure depiction highlighting the organisation of the helices (right). The bend-forming residues are in parentheses. Coiled structures are represented in white. **c**, Structural superimposition of the *Tb*MISP360 N-terminus crystal structure (deep purple) with a high confidence model of BARP (orange) and GARP (light blue, PDB: 2Y44) (left). A black dashed line highlights the area where the apical surface pocket is found. Top view of an electrostatic surface representation of *Tb*MISP360, BARP and GARP where the surface pocket is found (middle), and its orthogonal view indicating the residues forming the pocket and size (right). **d**, Conservation of the pocket-forming (top) and overall residues (bottom) across *Tb*MISP360, BARP and GARP depicted on the *Tb*MISP360 structure (left). Residue conservation between the five *T. brucei* MISP isoforms (right).

### A divergent surface pocket on MISP highlights the potential for ligand binding

Surface analysis of the *Tb*MISP360 and GARP^47^ crystal structures, and the high confidence homology model of BARP (generated here, see Methods), revealed a similar distribution of acidic and basic patches along the entire length of the structure with no clear localised charge densities that would indicate a molecular recognition site. However, a surface pocket was identified at the membrane distal end near the region where the core helices bend in both proteins (Fig. 5c). In *Tb*MISP360, the pocket is formed by the N-terminal portion of helix I, the loop connecting helix II to III, helix V and N-terminal region of helix VI. The secondary structures contributing to pocket formation in GARP were similar with the N-terminal portion of helix I, C-terminal region of helix II, the loop connecting helix III to IV, and N-terminal region of helix V forming the pocket. Intriguingly, the residues that form the pockets are quite divergent (Fig. 5c, right). For instance, the *Tb*MISP360 pocket (~12.2 Å in depth and 5.9 Å in diameter) is lined by ten hydrophobic residues (Ala^35^, Leu^37^, Phe^38^, Leu^42^, Ile^133^, Leu^169^ Ala^172^, Gly^176^, Ile^184^, and Leu^188^). In contrast, the pocket of GARP (~9.5 Å in depth and 6.4 Å in diameter) is formed by residues that include acidic, polar and hydrophobic (Thr^46^, Asp^49^, Val^50^, Gln^53^, Glu^168^, Thr^170^, Ser^171^, Ala^177^, Phe^178^, Leu^181^, Asp^190^). While the pocket in the BARP model (~10.1 Å in depth and 5.9 Å in diameter) also contains hydrophobic residues, it also contains several basic residues (Thr^36^, Ser^37^, Arg^38^, Ile^41^, Leu^44^, Arg^46^, Gln^50^, Arg^52^, Lys^140^, Leu^141^, Ala^173^, and Leu^174^, Arg^181^). The variability in the pocket composition prompted us to map the conserved residues between *Tb*MISP360, BARP and GARP on the *Tb*MISP360 structure using ConSurf ^48,49^ (Fig. 5d, left), which showed no conservation among the pocket-forming residues. Moreover, when mapping conservation between the five *T. brucei* MISP homologs on *Tb*MISP360, it was revealed that most of the conserved residues are found forming the core helices and the pocket core. However, a striking 51% of the non-conserved residues (which represent a 22.3% of the total sequence) localized right around the pocket (Fig. 5d, right), suggesting these residues may mediate affinity or recognition of the pocket’s putative ligand.

### Surface exposure of MISP suggests that MCFs express a more permissive VSG coat

A comparison between the structures of the N-termini of VSG MiTat 1.2 (PDB: 1VSG), MISP, and the BARP model (Supplementary Fig. 13a), revealed they are similar in height (~99.9 Å, ~93.0 Å and ~89.2 Å, respectively). However, when comparing the sequences of their divergent C-termini (Supplementary Fig. 13a), it was shown that *Tb*MISP360 and *Tb*MISP380 have ~71% and ~34% longer C-termini than VSGs; *Tb*MISP440 has an equivalent length and *Tb*MISP400/420 may have shorter (~60%) C-termini. Using the crystal structure of *Tb*MISP360 and the MISP C-terminal sequences, we generated structural models for all *T. brucei* MISP isoforms using I-TASSER^50-52^ and IntFOLD^53^ (Supplementary Fig. 13b). Like *Tc*HpHbR and VSG monomers, we predict that the MISP monomeric triple helical bundle is vertically oriented and, within the MCF surface context, is in close contact with neighboring mVSG homodimers. However, since MISP have longer C-terminal domains than VSGs, and despite their lack of secondary structure, we predict that the MISP N-termini (except for MISP400/420) are likely to extend above the mVSG coat (Fig. 6, Supplementary Fig. 13b). This is consistent with the MCF immunostaining, where the anti-MISP polyclonal antibody could specifically bind to MISP on live MCFs (Supplementary Fig. 8). Together with the results of immunostaining experiments with anti-BARP, these data suggest that the MCF surface is covered by a dense coat of mVSG homodimers, but it still allows room for the co-expression of invariant GPI-anchored glycoproteins, like BARP and MISP (the latter being more abundant and completely accessible to antibodies) (Fig. 6).

**Fig. 6.**
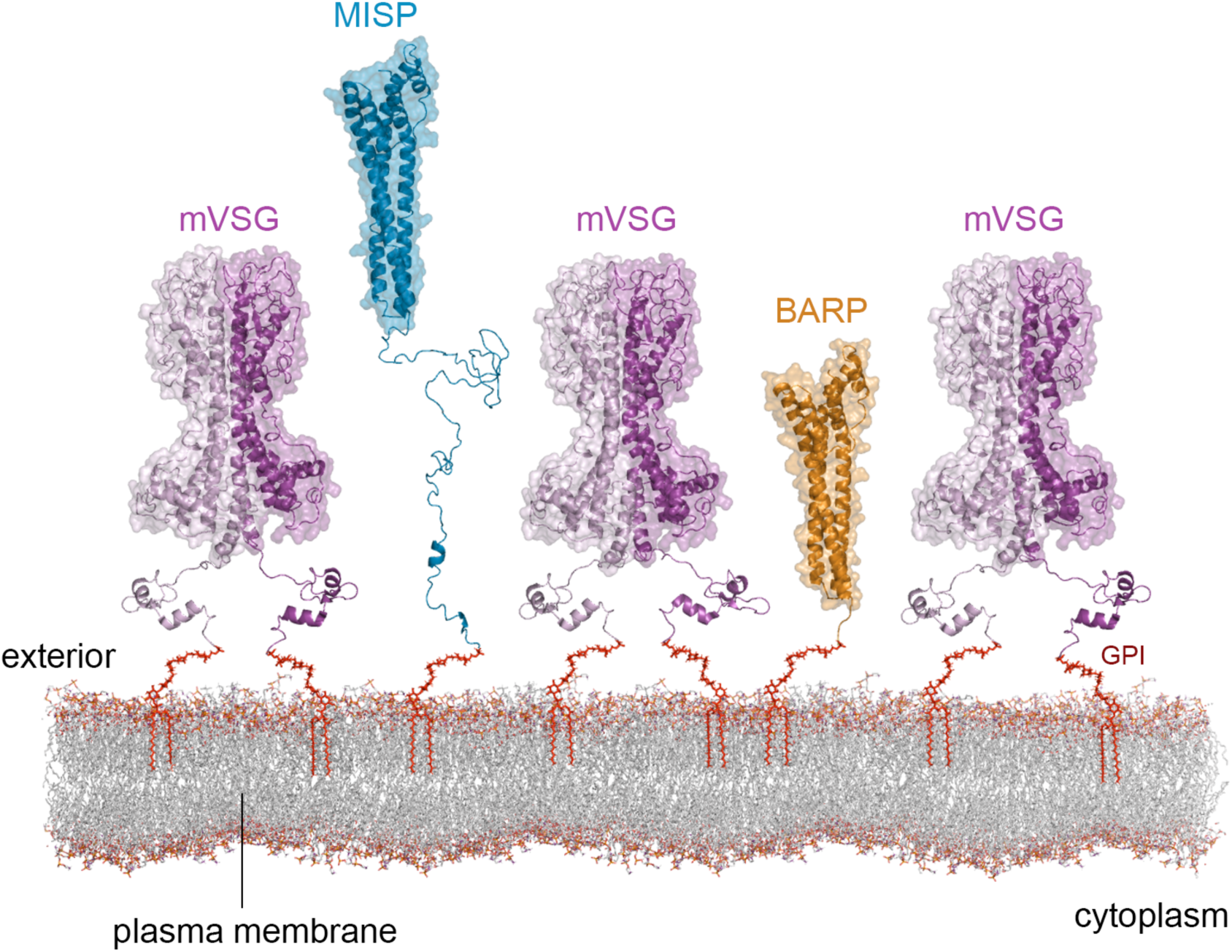
MISP and BARP define a novel metacyclic surface architecture. Structural model of the *T. brucei* metacyclic surface glycocalyx displaying the GPI-anchored metacyclic VSG homodimers (mVSG; light and deep blue), MISP (red) and remains of BARP (green). The mVSG structure is represented with a model of mVAT4 based on the crystal structure of VSG 221 N-(PDB: 1VSG) and C-terminus (PDB: 1XU6). MISP is represented with the crystal structure of *Tb*MISP360 N-terminus (PDB: 5VTL) and a model of its C-terminus; BARP is represented with a high confidence model.

### Are MISP important for *T. brucei* development within the tsetse salivary glands?

To gain more insight into the essentiality and function of MISP, we created a PCF mutant cell line (*misp^RNAi^)* to knockdown all *misp* isoforms through a tetracycline-inducible RNAi stem loop system^54^. As expected, based on the negligible transcript levels detected (Fig. 2a), silencing of *misp* in PCF did not show any growth or morphology phenotype *in vitro* (Fig. 7a). We then analyzed the essentiality of MISP by infecting tsetse with the *misp^RNAi^* cells in three biological replicates. Trypanosomes were isolated (30 dpi) from MGs, PVs, and SGs to determine infection prevalence and *misp* knockdown levels (Fig. 7b). In both PCF and MG procyclics, the transcript levels in Dox+ cells were non-significantly up (19%; p-value=0.04) and down regulated (5.9%; p-value=0.56), respectively, compared to the Dox-control. In contrast, when *misp* expression is naturally up regulated in PV and SG parasites (Fig. 2a), we observed a transcript reduction of 58.2% in PV (p-value=0.05) and 31.5% in SG trypanosomes (pooled samples from three replicates). Representative samples of MGs, PVs, and SGs were also processed for immunostaining to indirectly quantify MISP reduction using the anti-MISP polyclonal antibody. As in the wild type, all Dox+ or Dox-MG and PV forms were negative for staining. However, as described above, MCFs reacted with the antibody (Fig. 7c) and Dox+ MCFs showed a significant 63.1% reduction in mean fluorescence intensity (p-value <0.001) (Fig. 7d). No defects in cell shape or motility were observed. Furthermore, despite the reduction of MISP in Dox+ cells, no differences in tsetse MG, PV, and SG infection rates were observed compared to the Dox-control group (Fig. 7e).

**Fig. 7.**
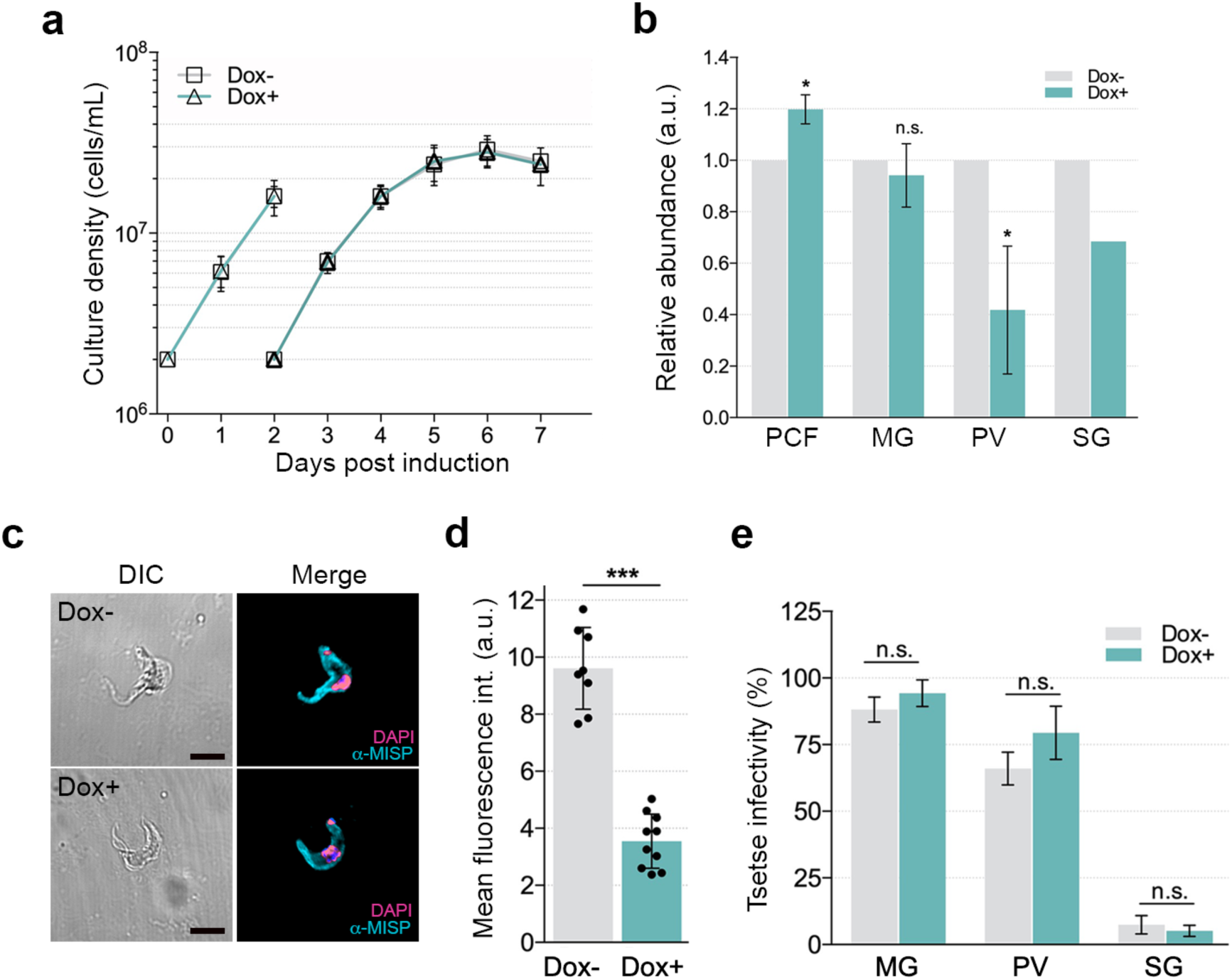
MISP do not appear to be essential for *T. b. brucei* development in the tsetse. **a**, *In vitro* growth curve of *misp^RNAi^* PCF cells, either induced (Dox+) or uninduced (Dox-). **b**, Relative *misp* mRNA expression in *misp^RNAi^* PCF cells, midgut procyclics (MG), proventricular parasites (PV) and salivary gland forms (SG). Expression levels of Dox-normalised to 1.0. **c**, Representative immunostaining images of non-permeabilised *misp^RNAi^* MCF (Dox-/+) with anti-MISP (red), DAPI (blue) and DIC. **d**, MISP mean fluorescence intensities (arbitrary units) on *misp^RNAi^* MCF (Dox-/+). **e**, Percentage of flies with *misp^RNAi^* cells infecting the midgut (MG), proventriculus (PV) and salivary glands (SG). Scale bars: 5 μm. Error bars show standard deviation, asterisks represent significance (* for p-value < 0.05; *** for p-value < 0.001).

## Discussion

### T. brucei-infected saliva is a complex mixture of tsetse, bacteria and trypanosome proteins

Transmission of vector-borne pathogens usually requires insect saliva as a “vehicle”, but it is also accompanied by a series of soluble components from both the parasite and the insect. For instance, in the sand fly, *Leishmania* spp. secrete abundant promastigote secretory gel and exosomes that are important virulence factors for transmission and establishment of the parasite infection in the vertebrate host^55–57^. We performed semi-quantitative proteomics to analyze the composition of *T. brucei-*infected tsetse saliva with the aim of identifying potential soluble factors involved in parasite transmission. Alongside hits from *Glossina*, we found proteins from the bacterial symbiont *S. glossinidius*, which compared to naïve flies seem to be more abundant in saliva during a trypanosome infection, as shown by immunoblotting. This could be explained in part by a higher cell permeabilization led by the trypanosome infection, as *Sodalis* is capable of intracellular and extracellular growth within tsetse, including the SGs^37^. An overall reduction in the amount of salivary proteins was confirmed in infected saliva as previously observed^32,34^. We also identified 27 different *T. brucei* proteins in infected saliva of which 62.9% are likely cytosolic and cytoskeletal. It remains unclear why trypanosome proteins are found in tsetse saliva and how they are released by the parasite. Their source could be dying parasites, and/or a more specific mechanism of protein secretion such as shedding of extracellular vesicles (EV)^58^. Indeed, it has been recently demonstrated that *T. brucei* BSFs secrete EVs that contain several virulence factors and proteins involved in parasite persistence within the host, including serum resistance-associated protein, VSGs, adenylate cyclase (GRESAG4), and GPI-PLC^59^. Notably, we showed that *T. brucei*-infected saliva is particularly enriched with trypanosome GPI-anchored surface proteins (37.1%); i.e. several VSGs, BARPs, and one (Tb927.7.360) belonging to a novel family of hypothetical proteins, herein named MISP, which we present as the first invariant surface proteins exposed on the surface of *T. brucei* MCF. Nevertheless, it remains to be determined whether MISP, VSGs, BARP, and other proteins here found in tsetse saliva are part of EVs, secreted in soluble form, or both.

### MISP are invariant surface proteins expressed by SG trypanosomes

We characterized the MISP family by studying gene and protein expression (during the parasite life cycle), cellular localization, essentiality for parasite development in the tsetse, and the crystal structure of the *Tb*MISP360 (Tb427.07.360) isoform. The majority of these studies were conducted alongside BARP as a known family of surface proteins expressed by EMFs in SGs^19^. In our proteomic analysis, we detected at least four different BARP isoforms in infected saliva (Tb927.9.15520, Tb927.9.15530 and Tb927.9.15570), in agreement with a previous report^34^. MISP were previously included in the trypanosome surface phylome^38^, within the clade ‘iv’ of the Fam50 family, which also includes BARP, and the *T. congolense* GARP^60,61^, and CESP^62^. Apart from *T. vivax*, most of the disease-relevant species of African trypanosomes such as *T. brucei* (including *T. b. gambiense* and *T. b. rhodesiense*)*, T. congolense,* and *T. evansi* encode *misp* genes. The preservation of MISP across these trypanosome species suggests a conserved function and an important role in the life cycle of these parasites.

We have confirmed that *misp* and *barp* gene expression is up regulated in *T. b. brucei* SG stages as previously determined^36,63,64,65^. However, our semi-quantitative RT-PCR method has allowed for the first time a detailed study of the individual expression of *misp* homologs by exploiting the different number of repetitive motifs at the C-termini. We found that *misp* up regulation is not homogeneous across homologs, but instead biased towards a preferential expression for *misp-B* paralogs (the shortest *misp400* and *misp420*). Except for *misp440* that seems to have a low but stable expression, *misp-A* transcripts are only detected during the late stages of parasite development in SG. It is unknown how these genes are expressed in such different ways despite being part of a gene array that is polycistronically-transcribed with >99.7% sequence conservation in UTR and intergenic sequences. Differences in homolog transcript levels, however, may not reflect changes in the respective protein abundance, which remain unknown as the polyclonal antibody used is expected to cross-react with all isoforms due to a high amino acid sequence conservation. Although PV trypanosomes already produce large amounts of *misp* and *barp* transcripts^19^ (Fig. 2), they are only translated in SG forms and localize evenly across the cell surface, as shown by immunostaining of individual SG forms. While BARP expression peaks in EMFs and is gradually reduced during metacyclogenesis, MISP expression begins in EMFs but reach their highest levels in MCFs, which led us to consider MISP as the predominant MCF protein. Strikingly, anti-MISP staining also occurs in live MCF cells (free of potential fixation artifacts), suggesting that MISP display immunogenic epitopes above the mVSG coat. Despite the previous observation that BARP is absent in MCFs^18,21^, we detected low levels of BARP in MCFs with variable localization, a phenomenon that could be explained by differences in parasite strain and/or dissection timing. While a minority of MCF cells showed a surface localization of BARP, most had re-distributed the protein to the flagellum. A similar effect was observed in BSFs^38^ and PCFs (Supplementary Fig. 9) where ectopic MISP initially localized to the flagella, but then regained surface location when cells differentiated to MCFs within the tsetse.

MCFs are cells with arrested cell cycle and low transcription activity, therefore detecting high levels of *misp* and *barp* transcripts suggests that these proteins could be abundant and play an important role in parasite development in the SG, transmission or early infection of the mammalian host. Given that RNAi silencing of *misp* in PV and SG forms did not seem to effect parasite viability or tsetse infectivity, MISP may rather play a role in transmission or early infection of the mammalian host^24^. However, we cannot rule out that MISP have a limited role in tsetse infectivity or that an insufficient knockdown did not lead to a detectable infection phenotype.

Having precisely defined that both MISP and BARP expression exclusively occurs in SG forms, our results allow a better understanding of the expression dynamics of major GPI-anchored surface glycoproteins during the life cycle of *T. brucei* (Fig. 8). In the blood stages, VSGs are clearly the dominant surface molecule^66^, which are then replaced by GPEET-followed by EP-procyclins after differentiation to the procyclic stage within the fly’s MG^15^. However, only EP-procyclins remain expressed in all PV parasite stages, including long and short EMFs (Supplementary Fig. 6). Once the parasites reach the tsetse SG, the procyclin coat is lost and BARP becomes the dominant molecule on attached EMFs^19^, while MISP gradually appear (Fig. 4). During metacyclogenesis, BARP are progressively replaced by MISP (Fig. 4) and mVSGs^22^ (Supplementary Fig. 10). Remarkably, some MCFs can also express BARP in low amounts and mostly in the flagellum (Fig. 4). The relative copy numbers of surface expressed BARP and MISP in relation to the mVSG coat remain unknown.

**Fig. 8.**
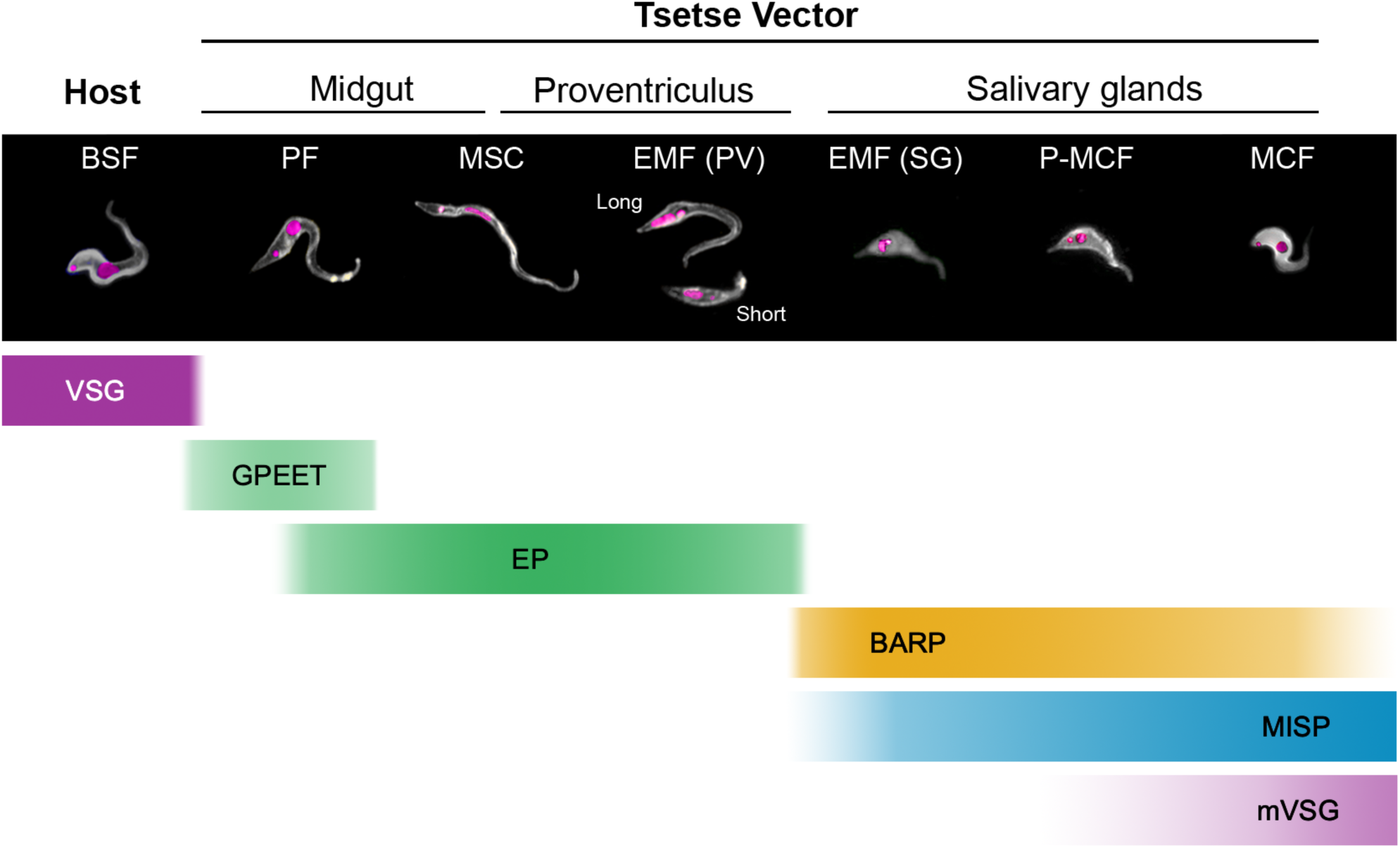
Developmental expression of the *T. b. brucei* major GPI-anchored surface glycoproteins during its life cycle. Bloodstream forms (BSF), procyclic forms (PF), mesocyclic forms (MSC), short and long epimastigotes infecting the proventriculus, attached epimastigotes infecting the salivary glands (EMF (SG)), pre-metacyclic forms (P-MCF) and metacyclic forms (MCF). Representative immunostaining images of the parasite stages highlighting the nuclei and kinetoplasts (magenta), and the cell surface (white) (top). Bottom bars define the duration and intensity of the expression of these proteins in relation to parasite developmental stage.

### The MISP helical bundle is exposed above the mVSG coat of *T. brucei* metacyclics

To gain more insight into the function of MISP, the crystal structure of the ordered N-terminal domain of the *Tb*MISP360 (Tb427.07.360) isoform was solved. It revealed a triple helical bundle structure that resembles other trypanosome surface proteins such as the *Tb/Tc*HpHbR^44,67^, GARP^47^ or a VSG monomer^46^. However, the mature *Tb*MISP360 contains an additional C-terminal tail predicted to be highly disordered. In the absence of definitive structural data, we modelled the C-terminus, which indicated that it might serve as a long tether effectively elevating the N-terminal head group from the surface of the parasite membrane. This explains (in part) the recognition of MISP by polyclonal antibodies shown in live immunostaining. It is tempting to speculate that this extension facilitates a biologically relevant interaction between the parasite and its environment such as promoting the acquisition of nutrients, serving as an adhesion or decoy, or modulating a response in the vertebrate host. The previously characterized *Tb*HpHbR lies partially within the VSG layer of the BSF and allows the trypanosome to acquire nutrients from the blood of the mammalian host. The mechanism by which it acquires nutrients is facilitated by a C-terminal extension that increases its height, thus making the ligand-binding site accessible by protruding above the VSG layer^68^. Since MCFs express mVSGs^69^ alongside MISP, a C-terminal extension could support a similar role although the substantial length of the extension would likely render cellular uptake of a nutrient payload difficult. However, the unstructured extension is also suited to support an adhesion role for MISP, similar to the function described for the *Haemophilus* Cha adhesins^70^. Intriguingly, the only study that defined an adhesin function of a trypanosome surface protein (CESP) is also part of the Fam50 family of proteins^62^, thereby suggesting that MISP could also play a role in adhesion. Our structural analyses revealed the existence of a predominantly hydrophobic pocket in the apical end of *Tb*MISP360 which may allow the coordination of a molecular partner, consistent with the putative role as an adhesin. Despite high sequence identity, the residues forming and surrounding the pocket lacked sequence conservation and resulted in a distorted cavity between the homolog models that may reflect the capacity to coordinate a structurally diverse set of ligands, potentially from different hosts (Supplementary Fig. 12). Furthermore, the homologs differ in the length of the C-terminal region, originating from the variable number of 26-residues motifs. This region could offer the parasite a robust mechanism to engage a variety of cell types or provide a sequential binding effect to enable tighter adhesion. It is noteworthy that these putative unstructured C-terminal regions contain lysine and arginine residues, which are inherently susceptible to proteolysis consistent with predictions using the PROSPER software^71^(Supplementary Fig. 13). The predicted proteolytic susceptibility may serve a strategic function in enabling the parasite to release the molecular interactions anchoring it to the tsetse salivary glands and thereby promote transmission.

### The surface of *T. brucei* MCF is more permissive to invariant surface proteins

It has been assumed that the MCF surface is structurally and functionally equivalent to that of BSFs as they show the same thick VSG (electron-dense) layer in TEM sections^11,22,72^. This implies that mVSGs also form a dense macromolecular barrier that possibly masks MCF invariant surface proteins from the host’s immune system. Most of the characterized BSF invariant surface proteins (e.g., ISG65, ISG75, ESAG4 and hexose transporters), which are found on either the flagellar or cell surface, are attached via transmembrane domains^7,10^. Two exceptions are the transferrin^8^ and the HpHbR receptors^9^ which are GPI-anchored and exclusively localize in the flagellar pocket. This restricted localization has been proposed to be due partly to the presence of only one GPI anchor molecule, differing from VSGs which form homodimers (i.e. two GPI anchors) that allow their homogeneous surface expression^73^. In contrast, our crystallographic and *in vivo* labelling evidence show that MISP appear to be displayed on the entire cell surface. This suggests that, unlike BSFs, MCFs allow the homogeneous surface expression of monomeric GPI-anchored invariant proteins. This clearly differs when MISP is ectopically expressed in BSF or PCF, in which the proteins localize at the flagellar pocket and/or the flagellar membrane. In summary, we hypothesize that MISP could be targeted by the vertebrate host’s immune system, although we do not know if the brief exposure of MISP in skin (directly following transmission) can trigger a protective and sustained immune response.

### Do metacyclic *vsg* genes recombine?

Our proteomic analysis identified peptides from several VSG species, which are responsible for antigenic variation in the blood stages. Although in the fly the parasite re-expresses VSGs only in the metacyclic stage, their role during development within the tsetse or during transmission remains unclear. The identification of mVSG peptides in infected saliva is not surprising given that infected SGs contain thousands of quiescent metacyclic cells, which may release mVSGs into the SG lumen. mVSG release may occur partly by the action of the parasite GPI-PLC, which is highly expressed in MCFs^21,74^ and/or by the action of tsetse salivary proteases. Interestingly, among the VSG species detected, only one corresponds to a canonical mVSG (mVAT4), which has not been previously detected as protein, but its MES has been well characterized^75^. Only a similar study detected mVAT5 in tsetse saliva infected, but from a different *T. brucei* strain^34^. Although it has been reported that BSFs may express mVSGs from BES at low levels^76^, no study has described the expression of BSF VSG proteins in MCFs. This supports the assumption that, although recombination may occur from MES to BES, it does not appear to occur in the opposite direction since MES lack the 70 bp repeats upstream of the mVSG gene. Surprisingly, however, we identified at least four BSF VSGs, including MiTat 1.2 (identified by three unique peptides). This suggests that either (1) MCF could have active BES (unlikely since MCFs do not appear to express ESAG proteins^74^) or 2) there is recombination at the MES replacing the *mvsg* by a *vsg* gene, or 3) there is recombination between *mvsg* and *vsg* genes leading to the formation of mosaics. In addition, we cannot rule out that the undetermined VSG repertoire of the TSW-196 strain^77^ (used in this study) could be different to that from other strains^78^. Importantly, previous studies have found similar results although this aspect has been overlooked^34,36,64,74^ (Supplementary Table 3). For example, at least two BSF VSGs were identified in saliva from flies infected with *T. b. brucei* EATRO 1125^34^; one BSF *vsg* was found to be highly transcribed in a *T. b. brucei* RUMP503 SG infection^64^; and 15 BSF *vsg*s had similar transcription levels than *mvsg*s (and a few were detected as protein) in *T. b. brucei* Lister 427 MCF overexpressing the RNA-binding protein 6^74^.

In summary, the *T. brucei-*infected tsetse saliva not only contains infective MCFs, but a cocktail of *G. morsitans* salivary proteins, bacterial and trypanosome soluble factors. How this complex composition modulates parasite transmission has yet to be understood. Among the detected parasite proteins in infected saliva, MISP represent the first family of immunogenic, invariant proteins described in *T. brucei*, which are exposed on the cell surface of the mammalian-infectious metacyclic stage. Based on their external exposure and sequence similarity, MISP could be ideal vaccine candidates against trypanosomiasis caused by *T. brucei* spp., in a similar way to the malaria vaccine that targets the sporozoite CSP protein^79^. Lastly, MISP have also the potential to be exploited as xenodiagnosis markers in the field for the identification of disease-transmissible tsetse.

## Methods

### Tsetse fly infection and dissection

*Glossina morsitans morsitans* were obtained from the tsetse insectary at the Liverpool School of Tropical Medicine, where they are maintained at 26 °C (+/-1 °C) and 65-75 % relative humidity. Flies were fed for 12 minutes every second day on sterile, defibrinated horse blood (TCS Biosciences) through a silicone membrane system. Teneral (24h post emergence, unfed) males were infected by combining *T. b. brucei* (strain TSW-196)-infected rat blood with fresh, sterile, defibrinated horse blood at a density of 10^6^ cells/mL. Alternatively, flies were infected by mixing *T. b. brucei* PCF cells (strain AnTat 1.1 90:13 or the derived mutant cell lines ^HA-eGFP^MISP360 and *misp*^RNAi^) with washed, defibrinated horse blood at a density of 5×10^5^ cells/ml. Uninfected control flies were fed on the same blood (free of trypanosomes) for the same amount of time. Any experimental flies that had not fed were removed the following day. Experimental flies were maintained for 4 weeks, feeding on normal defibrinated horse blood every 2 days. Alternatively, flies infected with mutant trypanosomes requiring tetracycline activation of the transgene were fed with normal defibrinated horse blood supplemented with fresh 2 μg/mL doxycycline (dissolved in 70% ethanol). Bloodmeals for control flies were supplemented with the same volume of 70% ethanol only. Flies were dissected at 28 to 30 days post infection, 72 hours after receiving the last bloodmeal. For saliva proteomics, based on normal *T. b. brucei* salivary gland infection rates (~15%) and yields (~10^4^ trypanosomes/gland pair), 1,500 naïve and ~1,500 *T. b. brucei*-infected tsetse were dissected. Salivary glands were collected in sterile ice-cold PBS and centrifuged at 12,000 rpm for 5 minutes at 4°C to collect the resultant saliva supernatant. SG and saliva were then separately stored at −80°C until further analyses. Alternatively, SG, PV and MG were dissected out (in this order to prevent trypanosome cross-contamination) in sterile ice-cold PBS and *T. brucei* cells were isolated, washed in PBS by centrifugation at 2,000 rpm for 5 minutes at 4°C and either stored at −80°C or directly used for immunostaining.

### *T. b. brucei* cell culture

*Trypanosoma brucei brucei* strain AnTat 1.1 90:13 PCF^54,80^ were cultured in SDM-79 medium supplemented with 10% FBS, haemin, 2 mM L-glutamine, Glutamax (Gibco), 15 μg/mL G418 and 50 μg/mL hygromycin B (Invitrogen), at 27°C and 5% CO_2_. Cells were typically passaged every 2 days. Mutant PCF cell lines were cultured in the presence of the selection marker drugs puromycin 1 μg/mL or zeocin 10 μg/mL (InvivoGen). Cells requiring the activation of the transgene were cultured in the presence of fresh 1μg/mL doxycycline (dissolved in 70% ethanol) or the same volume of 70% ethanol for negative controls. *T. b. brucei* AnTat 1.1 90:13 BSF were cultured in HMI-9 medium supplemented with 10% FBS, 10% Serum Plus and Glutamax (Gibco), 2.5 μg/mL G418 and 4 μg/mL hygromycin B at 37°C and 5% CO_2_.

### High-resolution nLC-MS/MS proteomic analysis

Samples from naïve and *T. brucei-*infected tsetse saliva were lyophilized and sent to the Dundee University Fingerprints Proteomics Facility for in-solution trypsin digests and identification by mass spectrometry. Following trypsinization, the resulting peptides were analyzed by nano-liquid chromatography tandem mass spectrometry (nLC-MS/MS) using the Ultimate 3000 RSLC nano coupled to LTQ-Orbitrap Velos Pro MS (Thermo Fisher Scientific). Resulting MS/MS spectra from HR-MS/MS analysis of *T. b. brucei* and *G. m. morsitans* samples were searched with Proteome Discoverer (PD) 2.1.1.21 (Thermo Fisher Scientific) and filtered with an estimated false-discovery rate (FDR) of 1%. The PD settings were as follows: HCD MS/MS; cysteine carboxyamidomethylation as fixed modification; methionine oxidation and acetylation on any amino acid as variable modifications; fully tryptic peptides only; up to two missed cleavages; parent ion mass tolerance of 10 ppm (monoisotopic); and fragment mass tolerance of 0.6 Da (in SEQUEST) and 0.02 Da (in PD 2.1.1.21) (monoisotopic). Tandem MS/MS spectra were searched against the following databases: a combined protein database of *T. b. brucei, T. b. gambiense* and *T. b. rhodesiense* (downloaded in FASTA format in October 2017 from TriTrypDB; http://www.tritrypdb.org/, 18,729 entries) and (downloaded in FASTA format in October 2017 from UniProtKB; http://www.uniprot.org/; 25,450 entries), and *G. m. morsitans* (downloaded in FASTA format in May 2017 from VectorBase; https://www.vectorbase.org/; 12,494 entries). Common contaminant sequences (MaxQuant) were included in this combined database. The resulting PD dataset was further processed through Scaffold Q+ 4.8.2 and perSPECtives (Proteome Software, Portland, OR). A protein threshold of 95%, peptide threshold of 95%, and a minimum number of one unique peptide was used for identification of proteins. Manual inspection of the MS/MS spectrum was always performed for identification of any protein based on a single peptide. Using BLAST2GO, proteins were BLAST searched against the non-redundant NCBI protein database and mapped and categorized according to gene ontology terms (GO). *Sodalis glossinidius* proteins were identified using the Liverpool School of Tropical Medicine in-house MASCOT MS/MS ion search using a protein database from UniProtKB generated from the latest re-annotated coding sequences in *S. glossinidius*^81^, using the following settings: two missed trypsin cleavages, same peptide modifications described above, peptide tolerance of 1.2 Da, MS/MS tolerance of 1.2 Da and peptide charges set to +1, +2 and +3.

### Bioinformatic analyses

All trypanosome sequences were obtained from TriTrypDB (www.tritrypdb.org) (Aslett, 2010 #133) or UniProt (www.uniprot.org)^82^. *G. m. morsitans* sequences were obtained from VectorBase (www.vectorbase.org)^83^. Sequence analyses and multiple alignments were done using Clustal Omega^84^ and Geneious R9 (Biomatters Ltd). Alignment figures were made in Jalview^85^. Phylogenetic trees were generated using PhyML 3.1^39^ under default conditions with a set bootstrap of 500. Signal peptide, GPI-anchor signal peptide, and transmembrane domain predictions were determined using the servers SignalP 4.1^40^, PredGPI^41^ and TMHMM 2.0^86,87^, respectively, under default settings. *N-*glycosylation predictions were run using the NetNGlyc 1.0 server (http://www.cbs.dtu.dk/services/NetNGlyc). MISP 3D protein modelling was performed using I-TASSER^50-52^ with the crystal structure of *Tb*MISP360 as a structural template and multiple sequence alignments (Clustal Omega) with the query sequences, or with IntFOLD^53^. Structural homology searches were performed using DALI^88^ and the conservation models using ConSurf^48,49^.

### Semi-quantitative RT-PCR

Total RNA from cultured and tsetse-isolated trypanosome samples was extracted using the GeneJet RNA extraction kit (Thermo Scientific) following the manufacturer’s protocol. Extracted RNA was further treated with the TURBO DNAse kit (Ambion) to remove residual genomic DNA following the manufacturer’s instructions. Variable amounts of RNA were reverse-transcribed into cDNA at 52°C for 10 minutes using SuperScript IV (Invitrogen) and ^oligo-dT20 primer, inactivated at 80°C for 10 minutes and treated with *E. coli* RNAse H at 37°C^ for 20 minutes. The resultant RNA-free cDNA was used in a downstream PCR using the Taq 2x Mastermix (New England Biolabs) and the primer pairs (400 nM) 5’-GCCGAAAGGACGGCAGAGACG-3’ and 5’-GTGTATCCTCCTATAGATT CTGCATAGC-3’ for the detection of individual *misp* homologs (amplicons of 410 bp, 332 bp, 254 bp and 176 bp for *misp360, misp380, misp440* and *misp400/420*, respectively); 5’-GGCTACTTTTGACAGTGTTATG-3’ and 5’-CTTCCTTCGCTTTCGTAACG-3’ to detect all *barp* homologs (270 bp amplicon). To determine *misp* knockdown levels in *misp^RNAi^* cells, the primers 5’-GATGCCTCTAAAGCGAGAGAAGG-3’ and 5’-GTACTGGCGGCACTATTGTCCTC-3’ were used (263 bp amplicon). Expression values were normalized against the expression of *tert*^89,90^ using the primers 5’-AACAGTAAGTCCCGTGTGG-3’ and 5’-CAAAGTCGTACTTCAACATCAG-3’ (233 bp amplicon). For all RT-PCR reactions, we ran a non-template control, a control for DNAse treatment, for reverse-transcription and a positive control with 1 ng genomic DNA for standardization. PCR conditions were as follows: 95° for 2 minutes, followed by 30 or 35 cycles of 95°C for 30 seconds, annealing for 30 seconds at variable temperatures, and elongation at 68°C for 30 seconds. A last elongation step at 68° was performed for 2 minutes. The cycle number was set at the exponential phase of the reaction. All RT-PCR products were mixed with sample buffer and run in a 2% agarose gel at 100V for 50 minutes. Gels were stained with SYBR Safe (Invitrogen), imaged using a Bio-Rad Gel Doc EZ system and analyzed using the ImageLab software. Imaging conditions were kept identical for all measurements for comparison.

### Immunofluorescence assay and confocal laser scanning microscopy

Cultured or tsetse-derived *T. b. brucei* cells were washed in PBS and fixed in fresh 4% paraformaldehyde (PFA) for 30 minutes at room temperature (RT). Cells were then allowed to settle onto poly-L-lysine coated slides for 20 minutes in a humidity chamber. Alternatively, cells were first air-dried on a poly-L-lysine slide, fixed in 100% methanol for 10 minutes at −20°C and rehydrated in PBS after air-drying. PFA-fixed cells were then either permeabilized or not in 0.1% Triton X-100 for 10 minutes at RT. They were then blocked in vPBS (PBS containing 45.9 mM sucrose and 10 mM glucose) 20% FBS for 1 hour, washed in PBS and incubated for 1 hour at RT with the primary antibody solution; 1:200 (anti-MISP, anti-BARP, anti-HA), 1:50 (anti-CRD), 1:100 (anti-PFR), 1:800 (anti-EP) dilutions in vPBS 20% FBS. After further washing, they were incubated for 1 hour at RT with the secondary antibody solution (1:500 goat-anti-mouse/rabbit Alexa Fluor 488/555; ThermoFisher). Cells were then incubated with 300 ng/mL 4',6-diamidino-2-phenylindole (DAPI) for 10 minutes, washed and mounted in Slowfade Diamond mounting oil (Life Technologies). Slides were imaged and analyzed using the confocal laser scanning microscope Zeiss LSM-880 and the Zeiss Zen software. For mean intensity fluorescence measurements, images were taken conserving laser power, pinhole aperture and digital gain settings. Images were further analyzed using Zen and ImageJ.

### Generation of ^HA-eGFP^MISP360 and *misp^RNAi^* PCF mutant cell lines

To create a cell line expressing ectopic MISP360 tagged with an HA-epitope and eGFP at N-terminus (^HA-eGFP^MISP360), the nucleotide sequence coding for the mature Tb427.07.360 protein lacking the signal peptide (Asp^18^ to Phe^382^) was obtained by PCR (Q5 high-fidelity, New England Biolabs) from *T. b. brucei* (strain AnTat 1.1 90:13) genomic DNA using the following primer sequences: 5’-GCG<u>TCTAGA</u>GACTCCATAATTGAGGAAGG-3’ and 5’-GCG<u>AGATCT</u>TTAAAAATGTGCGGCAGC-3’ with *XbaI* and *BglII* restriction enzyme targets (underlined), respectively. The Tb927.7.360 signal peptide (Met^1^ to Ala^17^) coding sequence was separately obtained by annealing the oligonucleotides 5’-CGC<u>AAGCTT</u>ATGACTAGCCCGTTTTTTGTGCTTGCTTGCTATACTTACGTATGTCACCGC C<u>AAGCTT</u>CGC-3’ and 5’-GCG<u>AAGCTT</u>GGCGGTGACATACGTAAGTATAGCAAG CAAGCACAAAAAACGGGCAGTCAT<u>AAGCTT</u>GCG-3’ with *HindIII* at both ends, and the HA epitope coding sequence was obtained by annealing the oligonucleotides 5’-CGC<u>AAGCTT</u>TATCCATATGACGTGCCGGATTATGCG<u>ACTAGT</u>CGC-3’ and 5’-GCG<u>ACTAGT</u>CGCATAATCCGGCACGTCATATGGATA<u>AAGCTT</u>GCG-3’ with *HindIII* and *SpeI*. The pDEX-577 plasmid^91^ for the tetracycline-inducible ectopic gene expression under a T7 promoter, was digested with *HindIII* and *SpeI* and ligated with *HindIII/SpeI*-digested HA sequence. The resultant plasmid was digested with *HindIII,* dephosphorylated with recombinant shrimp alkaline phosphatase and the *HindIII*-digested signal peptide coding sequence was inserted. The construct was then digested with *XbaI* and *BamHI* to ligate the *Tb927.7.360* (lacking the signal peptide) coding sequence digested with *XbaI* and *BglII*. The resultant construct was linearized using *NotI* high-fidelity (New England Biolabs) and transfected into *T. b. brucei* (strain AnTat 1.1 90:13) PCF using an Amaxa 4D nucleofector, program FI-115 in Tb-BSF buffer^92^. Clonal transfectant lines were selected with 10 μg/mL zeocin.

To create a cell line for the RNAi knockdown of *misp* (*misp^RNAi^*), a short unique 494 bp sequence highly conserved in all *misp* homologs was manually selected and obtained by PCR (Q5 high-fidelity NEB) using the primers 5’-CTACCACTACAGCAATTCGG-3’ and 5’- GCCTCTCGCAATTCGTCTTG-3’ with the restriction target pairs *XhoI/BamHI* or *NdeI/HindIII* respectively. By digesting with the corresponding restriction enzymes, the two amplicons were sequentially inserted in sense and antisense directions into the plasmid pALC14^54^ for a tetracycline-inducible transcription of a short hairpin double-stranded RNA. The plasmid was linearized (*NotI* high-fidelity) and transfected into AnTat 1.1 90:13 PCF as described above, and clonal transfectant cell lines were selected with 1 μg/mL puromycin. Transgene insertions were checked by PCR.

### Expression and purification of recombinant *Tb*MISP360 and BARP

The nucleotide sequence encoding *Tb*427.07.360 (*Tb*MISP360) (Gly^24^ to Ala^234^) and BARP ABB49055.1 (Glu^23^ to Gly^238^) were synthesized by GenScript and codon-optimized for *E. coli*. The constructs were sub-cloned into a modified pET32a vector (Novagen) containing N-terminal thioredoxin (Trx) and hexa-histidine (His_6_) tags separated from the gene of interest by a TEV protease site. Sequencing confirmed that no mutations were introduced. *E. coli* Rosetta-gami 2 (DE3) cells (Novagen) were transformed with the plasmid and grown in autoinduction medium (Invitrogen) from a 5% inoculum. Following four hours of growth at 37°C and 64 hours at 16°C, cells were harvested and the pellet re-suspended in 20 mM HEPES pH 8.3, 1 M NaCl, 30 mM imidazole and 5mM β-mercaptoethanol. Cells in suspension were lysed using a French Press, insoluble material was removed by centrifugation, and soluble fraction allowed to batch-bind with Ni-agarose beads at 4°C for 1 hour. Recombinant *Tb*MISP360 and BARP were eluted with the same buffer as before supplemented with 250 mM imidazole, and fractions were analyzed by SDS-PAGE and pooled based on purity. The Trx-His6 tag was removed by TEV cleavage overnight at 291 K, and the proteins were further purified by size exclusion chromatography (SEC) (Superdex 16/60 200) in HEPES buffered saline (HBS) with 1mM DTT. Protein concentration was determined by absorbance at 280 nm with calculated extinction coefficients of 10220 M^-1^ cm^-1^ (*Tb*MISP360), and 12740 M^-1^ cm^-1^ (BARP).

### Crystallization and data collection of *Tb*MISP360

Crystals of purified *Tb*MISP360 were initially identified in the Index screen (Molecular Dimensions) using the sitting drop method at 18°C. The final drops consisted of 0.3 μL of *Tb*MISP360 at 20 mg ml^-1^ with 0.3 μL of reservoir solution and were equilibrated against 60 μL of reservoir solution. Diffraction-quality crystals grew in 48 hours in 25% PEG, 1500. A single crystal was looped, cryopreserved in 12.5% glycerol for 20 seconds, and flash-cooled to 100 K directly in the cryostream. Diffraction data were collected on beamline 9-2 at Stanford Synchrotron Radiation Laboratory (SSRL) at a wavelength of 0.9795 Å.

### *Tb*MISP360 structure data processing, solution and refinement

Diffraction data to 1.82 Å resolution were processed using Imosflm^93^ and Scala^94^ in the CCP4 suite of programs^95^. Initial phases were obtained by molecular replacement using PHASER^96^ with one copy of GARP^47^ (PDB 2Y44; 15% identity over 210 residues). Solvent molecules were selected using COOT^97^ and refinement carried out using Phenix Refine^98^. The overall structure of *Tb*MISP360 was refined to an R_free_ of 20.43%. Stereo-chemical analysis performed with PROCHECK and SFCHECK in CCP4^95^ showed excellent stereochemistry with more than 99% of the residues in the favored conformations and no residues modelled in disallowed orientations of the Ramachandran plot. Overall 5% of the reflections were set aside for calculation of R_free_. Data collection and refinement statistics are presented in Supplementary Table 4. The atomic coordinates and structure factors have been deposited in the Protein Data Bank under the code 5VTL.

### *In silico* homology modelling of BARP

For modelling BARP, the crystal structure of GARP (PDB: 2Y44) and *Tb*MISP360 were used as the templates, with 24 and 28% identity, respectively. Using Modeller 9v18^99^, 10 models of BARP were generated and the best model was chosen based on the low value of normalised discrete optimized protein energy (DOPE). The assessment of the final model was carried out with Ramachandran statistics^100^, QMEAN^101^, and ProSA^102^.

### SDS-PAGE and Western Blotting

Trypanosome lysates were prepared by pelleting 10^6^ cells and heat-denaturing (98°C for 10 minutes) in Laemmli buffer. Samples were resolved on a 12.5% SDS-PAGE gel using the Bio-Rad mini-Protean system at 50V for 30 minutes followed by 2 hours at 120V. Proteins were transferred from the gel to a polyvinylidene fluoride (PVDF) membrane. After transfer at 90V for 1 hour, membranes were blocked (PBS-Tween 0.5 % v/v, 5 % w/v skim milk powder) for 1 hour at room temperature and incubated with primary anti-HA (1:10,000; Thermofisher) or anti-*Sodalis* Hsp60 mAb 1H1^103^ (mouse anti-symbiont GroEL; 1:10 dilution) overnight at 4 °C. The membranes were then washed in PBS-Tween (0.5% v/v) and probed at room temperature for 1 hour with secondary goat-anti-rabbit or goat-anti-mouse IgG horseradish peroxidase-conjugated antibodies (1:25,000 dilution; Thermofisher). The membranes were washed and probed with Super Signal West Dura substrate (Thermo Scientific) and the signal transferred onto X-ray Amersham Hyperfilm (GE Healthcare). Membranes were lastly stained in 0.2% (w/v) nigrosine in PBS for 15 minutes for loading control.

### Real-time RT-PCR

Quantitative real-time PCR was performed using an Agilent MX3005-P qPCR system. The reactions were set as follows in 20 μL reactions: 1X Luna Universal qPCR Mastermix (SYBR Green and ROX reference dye, New England Biolabs), 2 μL cDNA template and the primers (250 nM) 5’-GTGAGGAAGCAGAAGTTGG-3’ and 5’-AAAGTG CTGCAAGAAGGATC-3’ for *misp* detection and 5’-AAGCAAAGGTACAAGCAGAG-3’ and 5’-CGAGTGTTGCTCTCACAG-3’ for *barp* detection (amplicons of 84 bp and 96 bp, respectively). The *tert* gene was used as housekeeping gene for normalization, amplified using the primers 5’-AACAGTAAGTCCCGTGTGG-3’ and 5’-GCCTTCAGTT TGTCCAAGAAG-3’ with an expected amplicon of 80 bp. PCR thermal cycle conditions were as follows: 95°C for 1 minute, followed by 40 cycles of 95°C for 15 seconds and 60°C for 30 seconds. PCR product specificity was determined by final dissociation curve for all samples from 55°C to 95°C. Relative gene expression was calculated based on the cycle threshold (Ct)^85^ values applying the 2^-ΔΔCt^method^104^.

## Supporting information

## Acknowledgements

This work was partially supported by the Wellcome Trust project grant 093691/Z/10/Z (awarded to AAS), GlycoPar-EU FP7 Marie Curie Initial Training Network (GA. 608295) (awarded to ACS and AAS), MRC Concept in Confidence Award MC_PC_17167 (awarded to AAS) and the Natural Sciences and Engineering Research Council of Canada (awarded to MJB). ICA was partially supported by grant No. 2G12MD007592 from the National Institute of General Medical Sciences (NIGMS). We are grateful to Douglas Lamont (University of Dundee FingerPrints Proteomics Facility) for proteomics data and the Biomolecule Analysis Core Facility (BACF) at UTEP/BBRC, funded by NIGMS grant # 2G12MD007592, for part of the proteomic analyses. MJB gratefully acknowledges the Canada Research Chair program for salary support. The authors would like to thank the technical support staff at the Stanford Synchrotron Radiation Lightsource. Confocal imaging facilities at LSTM were funded by a Wellcome Trust Multi-User Equipment Grant (104936/Z/14/Z). We thank members of the Acosta-Serrano group for valuable input and, Dan Southern and Rob Leyland for maintaining the LSTM tsetse colony.

## Author contributions

ACS, SP, MJL, MJB, AAS conceived and designed experiments. ACS conducted molecular biology and microscopy work. ACS, SP, MAF, ICA, LRH, CY, and CR obtained and analyzed proteomics data. MJB and RR obtained MISP crystal structure and BARP modelling. ACS, MJB and AAS wrote the paper with input from all authors.

## Competing interests

The authors declare no competing financial interests.

## References

1 Giordani, F., Morrison, L. J., Rowan, T. G., HP, D. E. K. & Barrett, M. P. The animal trypanosomiases and their chemotherapy: a review. Parasitology 143, 1862–1889, doi:10.1017/S0031182016001268 (2016).

2 Matthews, K. R. The developmental cell biology of Trypanosoma brucei. J Cell Sci 118, 283–290, doi:10.1242/jcs.01649 (2005).

3 Horn, D. Antigenic variation in African trypanosomes. Mol Biochem Parasitol 195, 123–129, doi:10.1016/j.molbiopara.2014.05.001 (2014).

4 Mugnier, M. R., Stebbins, C. E. & Papavasiliou, F. N. Masters of Disguise: Antigenic Variation and the VSG Coat in Trypanosoma brucei. PLoS Pathog 12, e1005784, doi:10.1371/journal.ppat.1005784 (2016).

5 Pinger, J. et al. African trypanosomes evade immune clearance by O-glycosylation of the VSG surface coat. Nat Microbiol 3, 932–938, doi:10.1038/s41564-018-0187-6 (2018).

6 Engstler, M. et al. Hydrodynamic flow-mediated protein sorting on the cell surface of trypanosomes. Cell 131, 505–515, doi:10.1016/j.cell.2007.08.046 (2007).

7 Paindavoine, P. et al. A gene from the variant surface glycoprotein expression site encodes one of several transmembrane adenylate cyclases located on the flagellum of Trypanosoma brucei. Mol Cell Biol 12, 1218–1225 (1992).

8 Grab, D. J. et al. The transferrin receptor in African trypanosomes: identification, partial characterization and subcellular localization. Eur J Cell Biol 62, 114–126 (1993).

9 Vanhollebeke, B. et al. A haptoglobin-hemoglobin receptor conveys innate immunity to Trypanosoma brucei in humans. Science 320, 677–681, doi:10.1126/science.1156296 (2008).

10 Ziegelbauer, K., Multhaup, G. & Overath, P. Molecular characterization of two invariant surface glycoproteins specific for the bloodstream stage of Trypanosoma brucei. J Biol Chem 267, 10797–10803 (1992).

11 Vickerman, K. Developmental cycles and biology of pathogenic trypanosomes. Br Med Bull 41, 105–114 (1985).

12 Grandgenett, P. M., Otsu, K., Wilson, H. R., Wilson, M. E. & Donelson, J. E. A function for a specific zinc metalloprotease of African trypanosomes. PLoS Pathog 3, 1432–1445, doi:10.1371/journal.ppat.0030150 (2007).

13 Gruszynski, A. E., DeMaster, A., Hooper, N. M. & Bangs, J. D. Surface coat remodeling during differentiation of Trypanosoma brucei. J Biol Chem 278, 24665–24672, doi:10.1074/jbc.M301497200 (2003).

14 Aksoy, E. et al. Mammalian African trypanosome VSG coat enhances tsetse's vector competence. Proc Natl Acad Sci U S A 113, 6961–6966, doi:10.1073/pnas.1600304113 (2016).

15 Roditi, I. et al. Procyclin gene expression and loss of the variant surface glycoprotein during differentiation of Trypanosoma brucei. J Cell Biol 108, 737–746 (1989).

16 Acosta-Serrano, A. et al. The surface coat of procyclic Trypanosoma brucei: programmed expression and proteolytic cleavage of procyclin in the tsetse fly. Proc Natl Acad Sci U S A 98, 1513–1518, doi:10.1073/pnas.041611698 (2001).

17 Vassella, E. et al. A major surface glycoprotein of Trypanosoma brucei is expressed transiently during development and can be regulated post-transcriptionally by glycerol or hypoxia. Genes Dev 14, 615–626 (2000).

18 Sharma, R. et al. The heart of darkness: growth and form of Trypanosoma brucei in the tsetse fly. Trends Parasitol 25, 517–524, doi:10.1016/j.pt.2009.08.001 (2009).

19 Urwyler, S., Studer, E., Renggli, C. K. & Roditi, I. A family of stage-specific alanine-rich proteins on the surface of epimastigote forms of Trypanosoma brucei. Mol Microbiol 63, 218–228, doi:10.1111/j.1365-2958.2006.05492.x (2007).

20 Van Den Abbeele, J., Claes, Y., van Bockstaele, D., Le Ray, D. & Coosemans, M. Trypanosoma brucei spp. development in the tsetse fly: characterization of the post-mesocyclic stages in the foregut and proboscis. Parasitology 118 (Pt 5), 469–478 (1999).

21 Rotureau, B., Subota, I., Buisson, J. & Bastin, P. A new asymmetric division contributes to the continuous production of infective trypanosomes in the tsetse fly. Development 139, 1842–1850, doi:10.1242/dev.072611 (2012).

22 Tetley, L., Turner, C. M., Barry, J. D., Crowe, J. S. & Vickerman, K. Onset of expression of the variant surface glycoproteins of Trypanosoma brucei in the tsetse fly studied using immunoelectron microscopy. J Cell Sci 87 (Pt 2), 363–372 (1987).

23 Ginger, M. L. et al. Ex vivo and in vitro identification of a consensus promoter for VSG genes expressed by metacyclic-stage trypanosomes in the tsetse fly. Eukaryot Cell 1, 1000–1009 (2002).

24 Caljon, G. et al. The Dermis as a Delivery Site of Trypanosoma brucei for Tsetse Flies. PLoS Pathog 12, e1005744, doi:10.1371/journal.ppat.1005744 (2016).

25 Kolev, N. G., Ramey-Butler, K., Cross, G. A., Ullu, E. & Tschudi, C. Developmental progression to infectivity in Trypanosoma brucei triggered by an RNA-binding protein. Science 338, 1352–1353, doi:10.1126/science.1229641 (2012).

26 Hertz-Fowler, C. et al. Telomeric expression sites are highly conserved in Trypanosoma brucei. PLoS One 3, e3527, doi:10.1371/journal.pone.0003527 (2008).

27 Bangs, J.D. Evolution of Antigenic Variation in African Trypanosomes: Variant Surface Glycoprotein Expression, Structure, and Function. Bioessays, e1800181, doi:10.1002/bies.201800181 (2018).

28 Marcello, L. & Barry, J. D. Analysis of the VSG gene silent archive in Trypanosoma brucei reveals that mosaic gene expression is prominent in antigenic variation and is favored by archive substructure. Genome Res 17, 1344–1352, doi:10.1101/gr.6421207 (2007).

29 Turner, C. M., Barry, J. D., Maudlin, I. & Vickerman, K. An estimate of the size of the metacyclic variable antigen repertoire of Trypanosoma brucei rhodesiense. Parasitology 97 (Pt 2), 269–276 (1988).

30 Lenardo, M. J., Rice-Ficht, A. C., Kelly, G., Esser, K. M. & Donelson, J. E. Characterization of the genes specifying two metacyclic variable antigen types in Trypanosoma brucei rhodesiense. Proc Natl Acad Sci U S A 81, 6642–6646 (1984).

31 Jackson, D. G., Owen, M. J. & Voorheis, H. P. A new method for the rapid purification of both the membrane-bound and released forms of the variant surface glycoprotein from Trypanosoma brucei. Biochem J 230, 195–202 (1985).

32 Van Den Abbeele, J., Caljon, G., De Ridder, K., De Baetselier, P. & Coosemans, M. Trypanosoma brucei modifies the tsetse salivary composition, altering the fly feeding behavior that favors parasite transmission. PLoS Pathog 6, e1000926, doi:10.1371/journal.ppat.1000926 (2010).

33 Alves-Silva, J. et al. An insight into the sialome of Glossina morsitans morsitans. BMC Genomics 11, 213, doi:10.1186/1471-2164-11-213 (2010).

34 Kariithi, H. M., Boeren, S., Murungi, E. K., Vlak, J. M. & Abd-Alla, A. M. A proteomics approach reveals molecular manipulators of distinct cellular processes in the salivary glands of Glossina m. morsitans in response to Trypanosoma b. brucei infections. Parasit Vectors 9, 424, doi:10.1186/s13071-016-1714-z (2016).

35 Matetovici, I., Caljon, G. & Van Den Abbeele, J. Tsetse fly tolerance to T. brucei infection: transcriptome analysis of trypanosome-associated changes in the tsetse fly salivary gland. BMC Genomics 17, 971, doi:10.1186/s12864-016-3283-0 (2016).

36 Savage, A. F. et al. Transcriptome Profiling of Trypanosoma brucei Development in the Tsetse Fly Vector Glossina morsitans. PLoS One 11, e0168877, doi:10.1371/journal.pone.0168877 (2016).

37 Cheng, Q. & Aksoy, S. Tissue tropism, transmission and expression of foreign genes in vivo in midgut symbionts of tsetse flies. Insect Mol Biol 8, 125–132 (1999).

38 Jackson, A. P. et al. A cell-surface phylome for African trypanosomes. PLoS Negl Trop Dis 7, e2121, doi:10.1371/journal.pntd.0002121 (2013).

39 Guindon, S. & Gascuel, O. A simple, fast, and accurate algorithm to estimate large phylogenies by maximum likelihood. Syst Biol 52, 696–704 (2003).

40 Petersen, T. N., Brunak, S., von Heijne, G. & Nielsen, H. SignalP 4.0: discriminating signal peptides from transmembrane regions. Nat Methods 8, 785–786, doi:10.1038/nmeth.1701 (2011).

41 Pierleoni, A., Martelli, P. L. & Casadio, R. PredGPI: a GPI-anchor predictor. BMC Bioinformatics 9, 392, doi:10.1186/1471-2105-9-392 (2008).

42 Zamze, S. E., Ferguson, M. A., Collins, R., Dwek, R. A. & Rademacher, T. W. Characterization of the cross-reacting determinant (CRD) of the glycosyl-phosphatidylinositol membrane anchor of Trypanosoma brucei variant surface glycoprotein. Eur J Biochem 176, 527–534 (1988).

43 Holm, L. & Park, J. DaliLite workbench for protein structure comparison. Bioinformatics 16, 566–567 (2000).

44 Higgins, M. K. et al. Structure of the trypanosome haptoglobin-hemoglobin receptor and implications for nutrient uptake and innate immunity. Proc Natl Acad Sci U S A 110, 1905–1910, doi:10.1073/pnas.1214943110 (2013).

45 Stødkilde, K., Torvund-Jensen, M., Moestrup, S. K. & Andersen, C. B. Structural basis for trypanosomal haem acquisition and susceptibility to the host innate immune system. Nature communications 5 (2014).

46 Freymann, D. et al. 2.9 å resolution structure of the N-terminal domain of a variant surface glycoprotein from Trypanosoma brucei. Journal of molecular biology 216, 141–160 (1990).

47 Loveless, B. C. et al. Structural characterization and epitope mapping of the glutamic acid/alanine-rich protein from Trypanosoma congolense: defining assembly on the parasite cell surface. J Biol Chem 286, 20658–20665, doi:10.1074/jbc.M111.218941 (2011).

48 Ashkenazy, H. et al. ConSurf 2016: an improved methodology to estimate and visualize evolutionary conservation in macromolecules. Nucleic Acids Res 44, W344–350, doi:10.1093/nar/gkw408 (2016).

49 Ashkenazy, H., Erez, E., Martz, E., Pupko, T. & Ben-Tal, N. ConSurf 2010: calculating evolutionary conservation in sequence and structure of proteins and nucleic acids. Nucleic Acids Res 38, W529–533, doi:10.1093/nar/gkq399 (2010).

50 Roy, A., Kucukural, A. & Zhang, Y. I-TASSER: a unified platform for automated protein structure and function prediction. Nat Protoc 5, 725–738, doi:10.1038/nprot.2010.5 (2010).

51 Yang, J. et al. The I-TASSER Suite: protein structure and function prediction. Nat Methods 12, 7–8, doi:10.1038/nmeth.3213 (2015).

52 Zhang, Y. I-TASSER server for protein 3D structure prediction. BMC Bioinformatics 9, 40, doi:10.1186/1471-2105-9-40 (2008).

53 McGuffin, L. J., Atkins, J. D., Salehe, B. R., Shuid, A. N. & Roche, D. B. IntFOLD: an integrated server for modelling protein structures and functions from amino acid sequences. Nucleic Acids Res 43, W169–173, doi:10.1093/nar/gkv236 (2015).

54 MacGregor, P., Rojas, F., Dean, S. & Matthews, K. R. Stable transformation of pleomorphic bloodstream form Trypanosoma brucei. Mol Biochem Parasitol 190, 60–62, doi:10.1016/j.molbiopara.2013.06.007 (2013).

55 Rogers, M. E. The role of Leishmania proteophosphoglycans in sand fly transmission and infection of the Mammalian host. Front Microbiol 3, 223, doi:10.3389/fmicb.2012.00223 (2012).

56 Atayde, V. D. et al. Exosome Secretion by the Parasitic Protozoan Leishmania within the Sand Fly Midgut. Cell Rep 13, 957–967, doi:10.1016/j.celrep.2015.09.058 (2015).

57 Giraud, E. et al. Leishmania proteophosphoglycans regurgitated from infected sand flies accelerate dermal wound repair and exacerbate leishmaniasis via insulin-like growth factor 1-dependent signalling. PLoS Pathog 14, e1006794, doi:10.1371/journal.ppat.1006794 (2018).

58 Colombo, M., Raposo, G. & Thery, C. Biogenesis, secretion, and intercellular interactions of exosomes and other extracellular vesicles. Annual review of cell and developmental biology 30, 255–289, doi:10.1146/annurev-cellbio-101512-122326 (2014).

59 Szempruch, A. J. et al. Extracellular Vesicles from Trypanosoma brucei Mediate Virulence Factor Transfer and Cause Host Anemia. Cell 164, 246–257, doi:10.1016/j.cell.2015.11.051 (2016).

60 Beecroft, R. P., Roditi, I. & Pearson, T. W. Identification and characterization of an acidic major surface glycoprotein from procyclic stage Trypanosoma congolense. Mol Biochem Parasitol 61, 285–294 (1993).

61 Bayne, R. A., Kilbride, E. A., Lainson, F. A., Tetley, L. & Barry, J. D. A major surface antigen of procyclic stage Trypanosoma congolense. Mol Biochem Parasitol 61, 295–310 (1993).

62 Sakurai, T., Sugimoto, C. & Inoue, N. Identification and molecular characterization of a novel stage-specific surface protein of Trypanosoma congolense epimastigotes. Mol Biochem Parasitol 161, 1–11, doi:10.1016/j.molbiopara.2008.05.003 (2008).

63 Savage, A. F. et al. Transcript expression analysis of putative Trypanosoma brucei GPI-anchored surface proteins during development in the tsetse and mammalian hosts. PLoS Negl Trop Dis 6, e1708, doi:10.1371/journal.pntd.0001708 (2012).

64 Telleria, E. L. et al. Insights into the trypanosome-host interactions revealed through transcriptomic analysis of parasitized tsetse fly salivary glands. PLoS Negl Trop Dis 8, e2649, doi:10.1371/journal.pntd.0002649 (2014).

65 Awuoche, E. O. et al. Expression profiling of Trypanosoma congolense genes during development in the tsetse fly vector Glossina morsitans morsitans. Parasit Vectors 11, 380, doi:10.1186/s13071-018-2964-8 (2018).

66 Cross, G. A. Identification, purification and properties of clone-specific glycoprotein antigens constituting the surface coat of Trypanosoma brucei. Parasitology 71, 393–417 (1975).

67 Lane-Serff, H., MacGregor, P., Lowe, E. D., Carrington, M. & Higgins, M. K. Structural basis for ligand and innate immunity factor uptake by the trypanosome haptoglobin-haemoglobin receptor. Elife 3, e05553, doi:10.7554/eLife.05553 (2014).

68 Higgins, M. K. et al. Structure of the trypanosome haptoglobin–hemoglobin receptor and implications for nutrient uptake and innate immunity. Proceedings of the National Academy of Sciences 110, 1905–1910 (2013).

69 Graham, S., Matthews, K., Shiels, P. & Barry, J. Distinct, developmental stage-specific activation mechanisms of trypanosome VSG genes. Parasitology 101, 361–367 (1990).

70 McCann, J. R., Sheets, A. J., Grass, S. & St Geme, J. W., 3rd. The Haemophilus cryptic genospecies Cha adhesin has at least two variants that differ in host cell binding, bacterial aggregation, and biofilm formation properties. J Bacteriol 196, 1780–1788, doi:10.1128/JB.01409-13 (2014).

71 Song, J. et al. PROSPER: an integrated feature-based tool for predicting protease substrate cleavage sites. (2012).

72 Vickerman, K. On the surface coat and flagellar adhesion in trypanosomes. J Cell Sci 5, 163–193 (1969).

73 Tiengwe, C., Bush, P. J. & Bangs, J. D. Controlling transferrin receptor trafficking with GPI-valence in bloodstream stage African trypanosomes. PLoS Pathog 13, e1006366, doi:10.1371/journal.ppat.1006366 (2017).

74 Christiano, R. et al. The proteome and transcriptome of the infectious metacyclic form of Trypanosoma brucei define quiescent cells primed for mammalian invasion. Mol Microbiol, doi:10.1111/mmi.13754 (2017).

75 Pedram, M. & Donelson, J. E. The anatomy and transcription of a monocistronic expression site for a metacyclic variant surface glycoprotein gene in Trypanosoma brucei. J Biol Chem 274, 16876–16883 (1999).

76 Alarcon, C. M., Son, H. J., Hall, T. & Donelson, J. E. A monocistronic transcript for a trypanosome variant surface glycoprotein. Mol Cell Biol 14, 5579–5591 (1994).

77 Paindavoine, P. et al. The use of DNA hybridization and numerical taxonomy in determining relationships between Trypanosoma brucei stocks and subspecies. Parasitology 92 (Pt 1), 31–50 (1986).

78 Cross, G. A., Kim, H. S. & Wickstead, B. Capturing the variant surface glycoprotein repertoire (the VSGnome) of Trypanosoma brucei Lister 427. Mol Biochem Parasitol 195, 59–73, doi:10.1016/j.molbiopara.2014.06.004 (2014).

79 Gordon, D. M. et al. Safety, immunogenicity, and efficacy of a recombinantly produced Plasmodium falciparum circumsporozoite protein-hepatitis B surface antigen subunit vaccine. J Infect Dis 171, 1576–1585 (1995).

80 Engstler, M. & Boshart, M. Cold shock and regulation of surface protein trafficking convey sensitization to inducers of stage differentiation in Trypanosoma brucei. Genes Dev 18, 2798–2811, doi:10.1101/gad.323404 (2004).

81 Belda, E., Moya, A., Bentley, S. & Silva, F. J. Mobile genetic element proliferation and gene inactivation impact over the genome structure and metabolic capabilities of Sodalis glossinidius, the secondary endosymbiont of tsetse flies. BMC Genomics 11, 449, doi:10.1186/1471-2164-11-449 (2010).

82 Apweiler, R. et al. The InterPro database, an integrated documentation resource for protein families, domains and functional sites. Nucleic Acids Res 29, 37–40 (2001).

83 Lawson, D. et al. VectorBase: a home for invertebrate vectors of human pathogens. Nucleic Acids Res 35, D503–505, doi:10.1093/nar/gkl960 (2007).

84 Sievers, F. et al. Fast, scalable generation of high-quality protein multiple sequence alignments using Clustal Omega. Mol Syst Biol 7, 539, doi:10.1038/msb.2011.75 (2011).

85 Waterhouse, A. M., Procter, J. B., Martin, D. M., Clamp, M. & Barton, G. J. Jalview Version 2−-a multiple sequence alignment editor and analysis workbench. Bioinformatics 25, 1189–1191, doi:10.1093/bioinformatics/btp033 (2009).

86 Sonnhammer, E. L., von Heijne, G. & Krogh, A. A hidden Markov model for predicting transmembrane helices in protein sequences. Proc Int Conf Intell Syst Mol Biol 6, 175–182 (1998).

87 Krogh, A., Larsson, B., von Heijne, G. & Sonnhammer, E. L. Predicting transmembrane protein topology with a hidden Markov model: application to complete genomes. J Mol Biol 305, 567–580, doi:10.1006/jmbi.2000.4315 (2001).

88 Holm, L. & Rosenstrom, P. Dali server: conservation mapping in 3D. Nucleic Acids Res 38, W545–549, doi:10.1093/nar/gkq366 (2010).

89 Brenndorfer, M. & Boshart, M. Selection of reference genes for mRNA quantification in Trypanosoma brucei. Mol Biochem Parasitol 172, 52–55, doi:10.1016/j.molbiopara.2010.03.007 (2010).

90 Dewar, S. et al. The Role of Folate Transport in Antifolate Drug Action in Trypanosoma brucei. J Biol Chem 291, 24768–24778, doi:10.1074/jbc.M116.750422 (2016).

91 Kelly, S. et al. Functional genomics in Trypanosoma brucei: a collection of vectors for the expression of tagged proteins from endogenous and ectopic gene loci. Mol Biochem Parasitol 154, 103–109, doi:10.1016/j.molbiopara.2007.03.012 (2007).

92 Burkard, G., Fragoso, C. M. & Roditi, I. Highly efficient stable transformation of bloodstream forms of Trypanosoma brucei. Mol Biochem Parasitol 153, 220–223, doi:10.1016/j.molbiopara.2007.02.008 (2007).

93 Leslie, A. G. W. Recent changes to the MOSFLM package for processing film and image plate data. Joint CCP4 + ESF-EAMCB Newsletter on Protein Crystallography 26 (1992).

94 Evans, P. R. Scaling and assessment of data quality. Acta Cryst D, 72–82 (2005).

95 Collaborative Computational Project, N. Acta Crystallogr. Sect. D Biol. Crystallogr. 50, 760–763 (1994).

96 McCoy, A. J. et al. Phaser crystallographic software. J Appl Crystallogr 40, 658–674, doi:10.1107/S0021889807021206 (2007).

97 Emsley, P. & Cowtan, K. Coot: model-building tools for molecular graphics. Acta Crystallog. sect. D 60, 2126–2132 (2004).

98 Adams, P. D. et al. PHENIX: a comprehensive Python-based system for macromolecular structure solution. Acta Crystallogr D Biol Crystallogr 66, 213–221, doi:10.1107/S0907444909052925 (2010).

99 Webb, B., Sali, A. Protein Structure Modelling with MODELLER. Functional Genomics, 39–54 (2016).

100 Lovell, S. C. et al. Structure validation by Calpha geometry: phi,psi and Cbeta deviation. Proteins 50, 437–450, doi:10.1002/prot.10286 (2003).

101 Benkert, P., Tosatto, S. C. & Schomburg, D. QMEAN: A comprehensive scoring function for model quality assessment. Proteins 71, 261–277, doi:10.1002/prot.21715 (2008).

102 Wiederstein, M. & Sippl, M. J. ProSA-web: interactive web service for the recognition of errors in three-dimensional structures of proteins. Nucleic Acids Res 35, W407-410, doi:10.1093/nar/gkm290 (2007).

103 Rose, C. et al. An investigation into the protein composition of the teneral Glossina morsitans morsitans peritrophic matrix. PLoS Negl Trop Dis 8, e2691, doi:10.1371/journal.pntd.0002691 (2014).

104 Livak, K. J. & Schmittgen, T. D. Analysis of relative gene expression data using real-time quantitative PCR and the 2(-Delta Delta C(T)) Method. Methods 25, 402–408, doi:10.1006/meth.2001.1262 (2001).

