## Supplementary material for "The crystal structure and localization of *Trypanosoma brucei* invariant surface glycoproteins suggest a more permissive VSG coat in the tsetse-transmitted metacyclic stage"

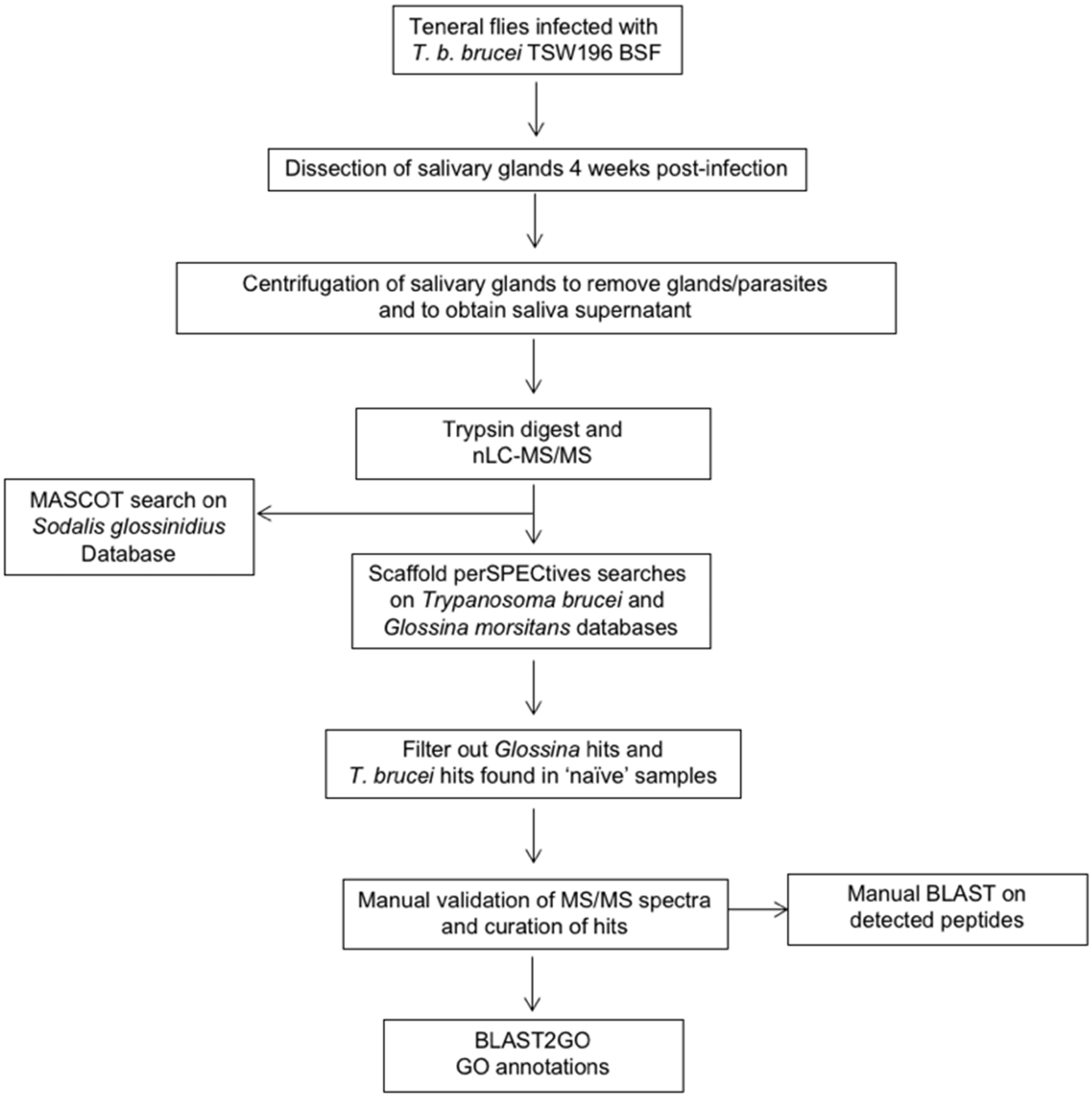

**Supplementary Fig. 1 | Workflow for the analysis of *T. b.* brucei-infected tsetse saliva.** For each biological replicate, saliva samples from naïve flies were also collected and processed in parallel.

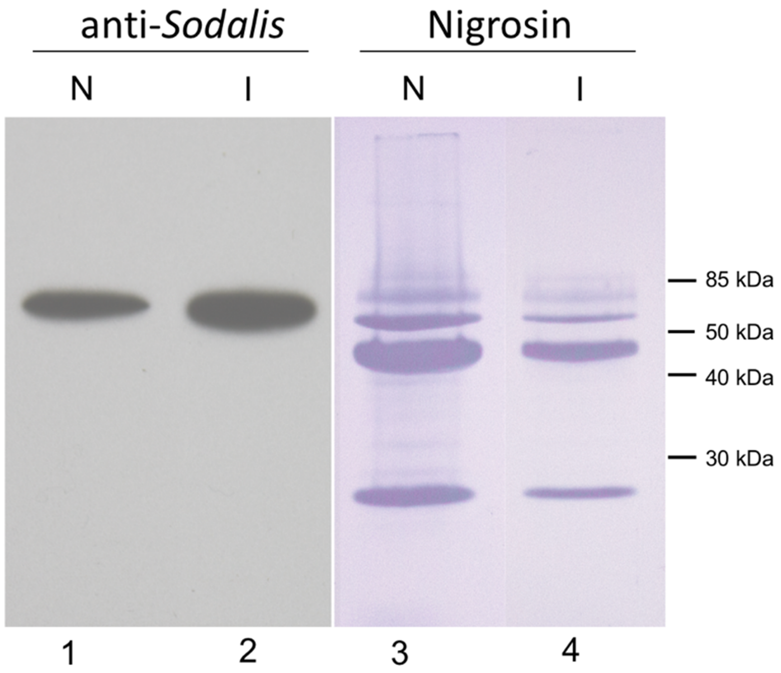

**Supplementary Fig. 2 | Immunoblotting of *G. m. morsitans* naïve and *T. b. brucei*-infected saliva probing with the mouse anti-*Sodalis* mAb 1H1 (recognizes Hsp60 from *Sodalis glossinidius*).** (Lane 1) Naïve saliva (N) and (Lane 2) infected saliva (I) probed with the antibody; film exposure: 10 seconds. Lanes 3 and 4 show the same PVDF membrane stained with nigrosine as loading control.

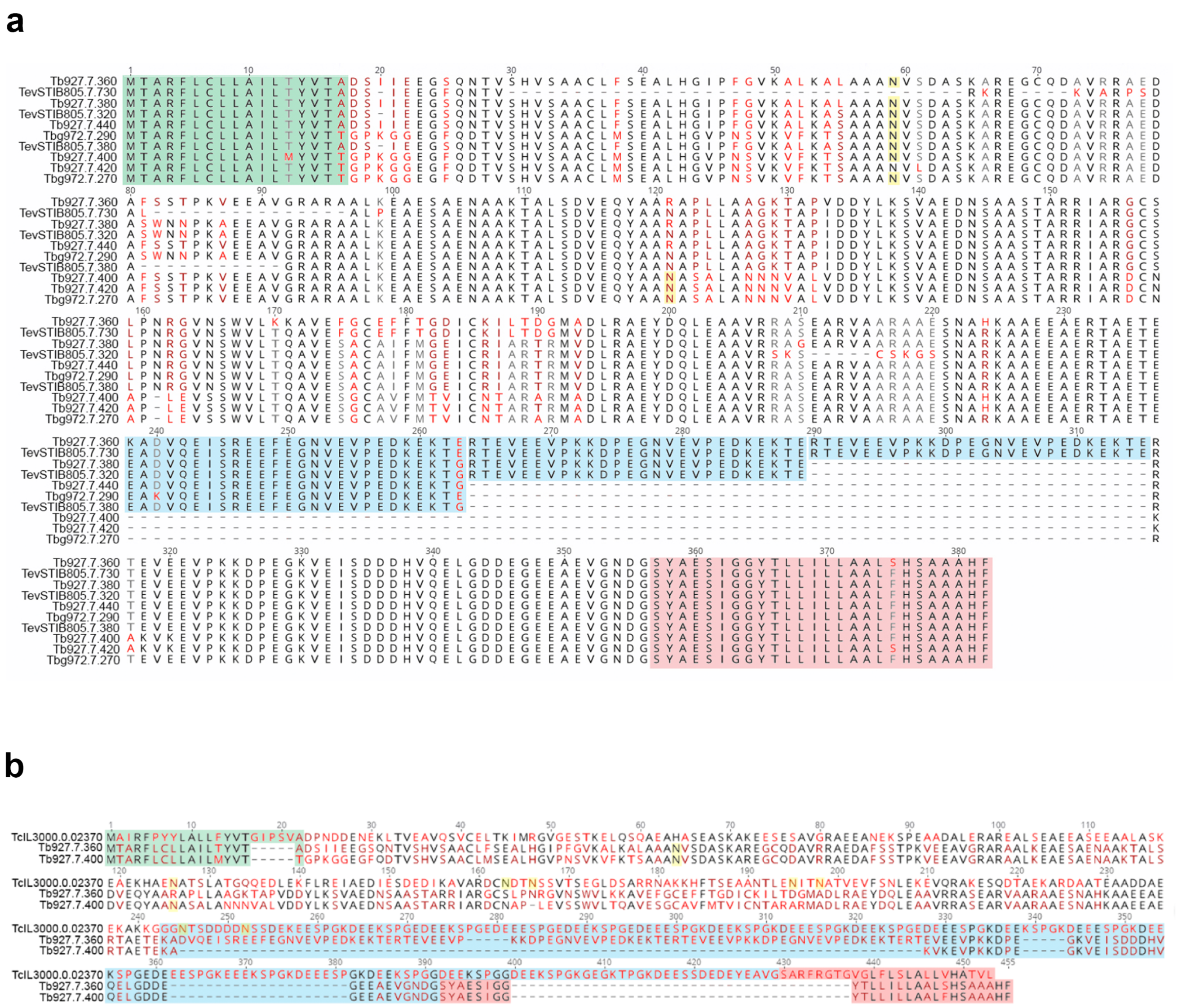

**Supplementary Fig. 3 | Multiple protein sequence alignments of MISP. a**, Multiple protein alignment of all MISP (*T. congolense* MISP excluded). Residues are coloured based on identity conservation from black (maximum) to red (minimum). Alignment made using a Clustal Omega BLOSUM62 matrix with a gap open cost 10 and a gap extension cost of 0.1, using Geneious R9. Domains highlighted: signal peptide (green), predicted *N*-glycosylation sites (yellow asparagines), C-terminal motifs (blue), and GPI-anchor attachment signal peptide (red). **b**, Multiple protein alignment of *T. congolense* MISP with *Tb*MISP360 (MISP-A) and *Tb*MISP400 (MISP-B). Domains highlighted as in 'a'.

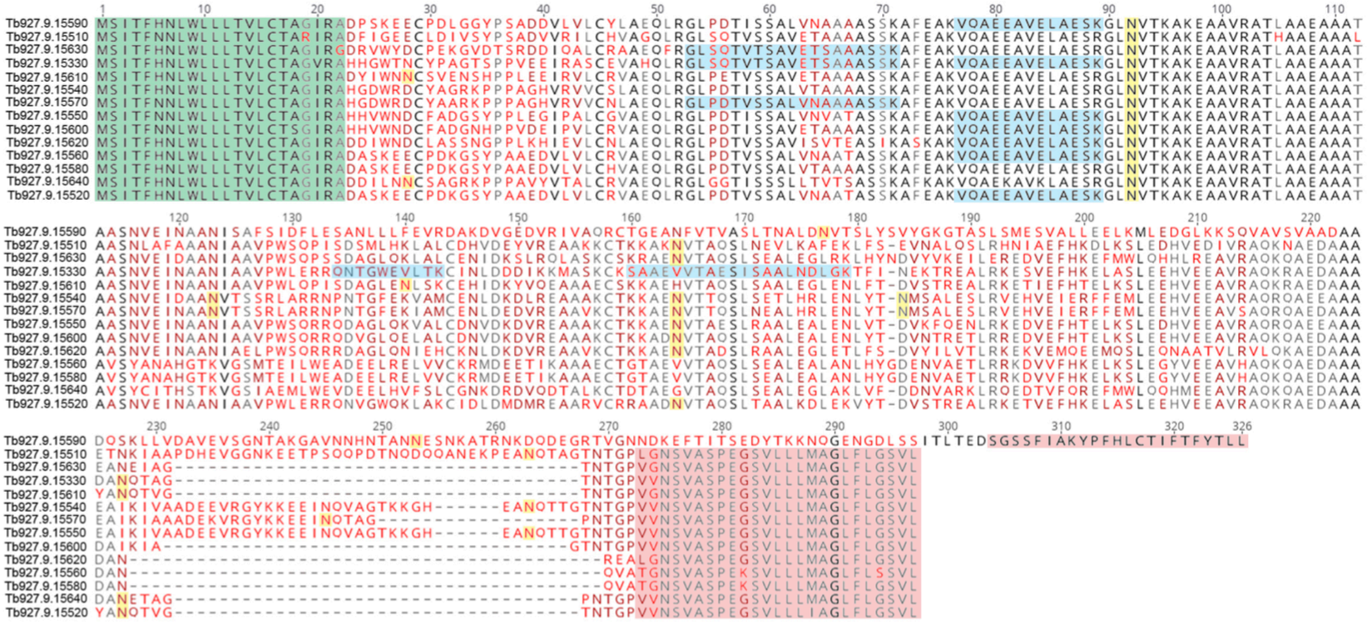

**Supplementary Fig. 4 | Multiple protein alignment of *T. b. brucei* BARP.** Residue colours are based on identity conservation from black (maximum) to red (minimum). Alignment made using a Clustal Omega BLOSUM62 matrix with a gap open cost 10 and a gap extension cost of 0.1, using Geneious R9. The BARP peptides detected in the nLC-MS/MS analysis are highlighted in blue. Domains highlighted: signal peptides (green), predicted *N*-glycosylation sites (yellow asparagines) and GPI-anchor attachment signal peptides (red).

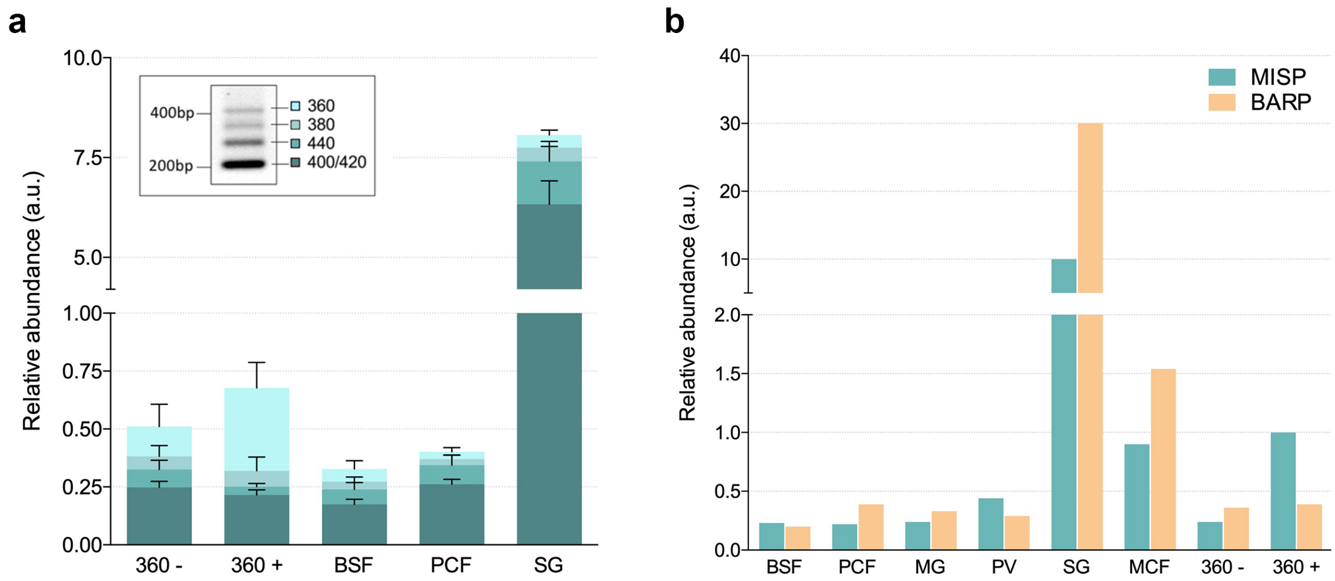

**Supplementary Fig. 5 | Relative expression of *misp* homologs and *barp*. a**, Controls for the semi-quantitative RT-PCR method used to determine the relative *misp* mRNA expression levels. BSF, PCF and SG samples shown as in Fig. 2. $^{HA\text{-}eGFP}$MISP360 PCF cells were introduced as controls, either uninduced (360-) or induced (360+). The inset is a representative image of a DNA agarose gel displaying 4 bands corresponding to the RT-PCR amplification of the different *misp* homologs. Expression normalised to the expression of *tert*. **b**, *misp* and *barp* relative RNA expression levels using quantitative RT-PCR on the samples in Fig. 2, including the controls used above.

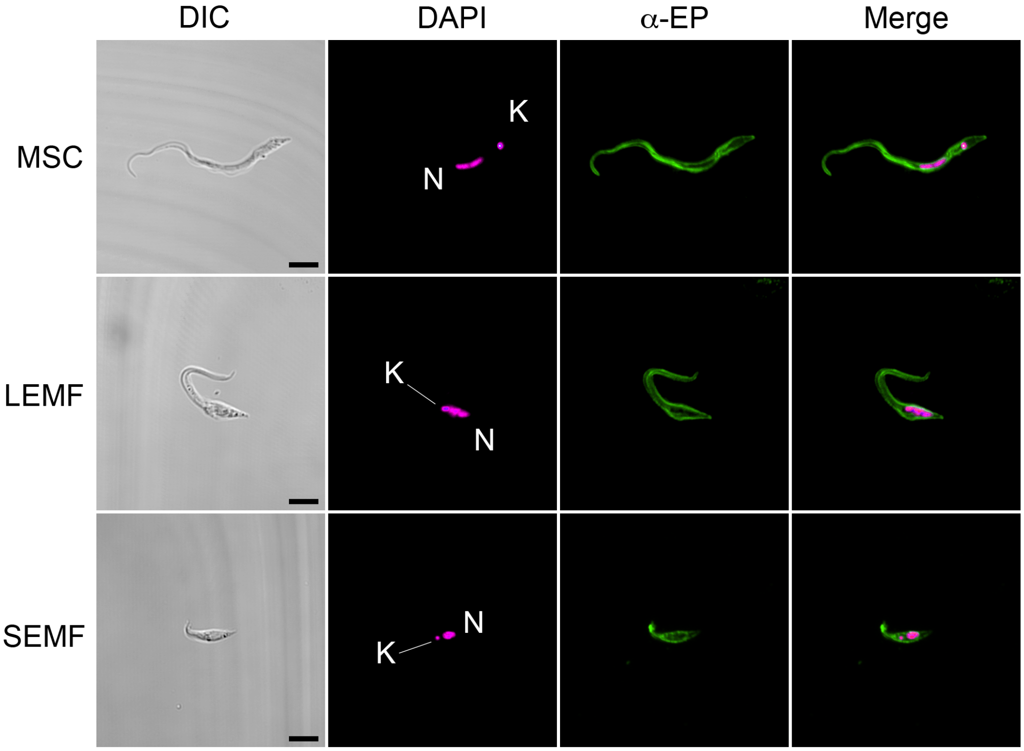

**Supplementary Fig. 6 | Expression of EP-procyclins in proventricular trypanosomes.** Parasites extracted from infected PV at 30 dpi. Mesocyclics (MSC), long epimastigote forms (LEMF) and short epimastigote forms (SEMF) shown by differential interference contrast (DIC) and probed with anti-EP monoclonal antibody (yellow) and DAPI (blue). Nuclei (N) and kinetoplasts (K) noted in the blue channel. Scale bars: 5 μm.

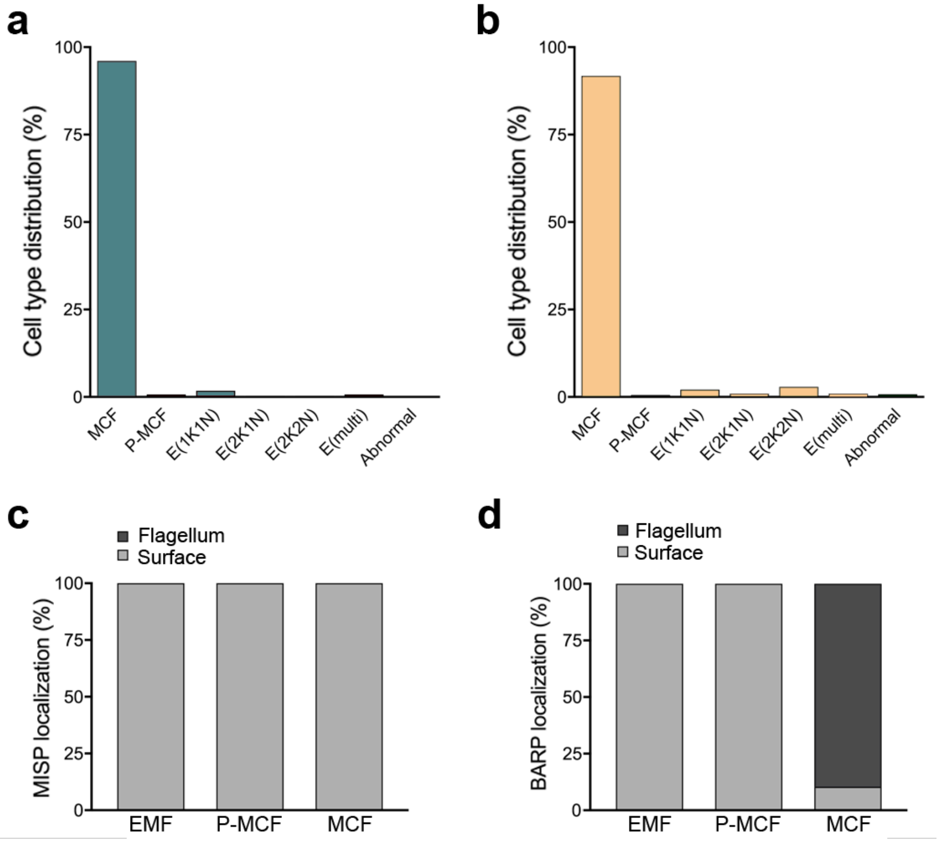

**Supplementary Fig. 7 | Trypanosome population in the tsetse SG.** Cell type distribution (percentage) in SG-extracted parasites used for the quantification of mean fluorescence intensities of MISP (**a**) and BARP (**b**). Metacyclic forms (MCF), pre-metacyclic forms (P-MCF), epimastigotes with variable kDNA (K) and nuclei (N) numbers (E(1K1N), E(2K1N), E(2K2N)), multi-nucleated epimastigotes (E(multi)) and abnormal cells. Distribution (percentage) of localisation types for MISP (**c**) and BARP (**d**), either flagellar (flagellum) or on the whole cell surface (surface).

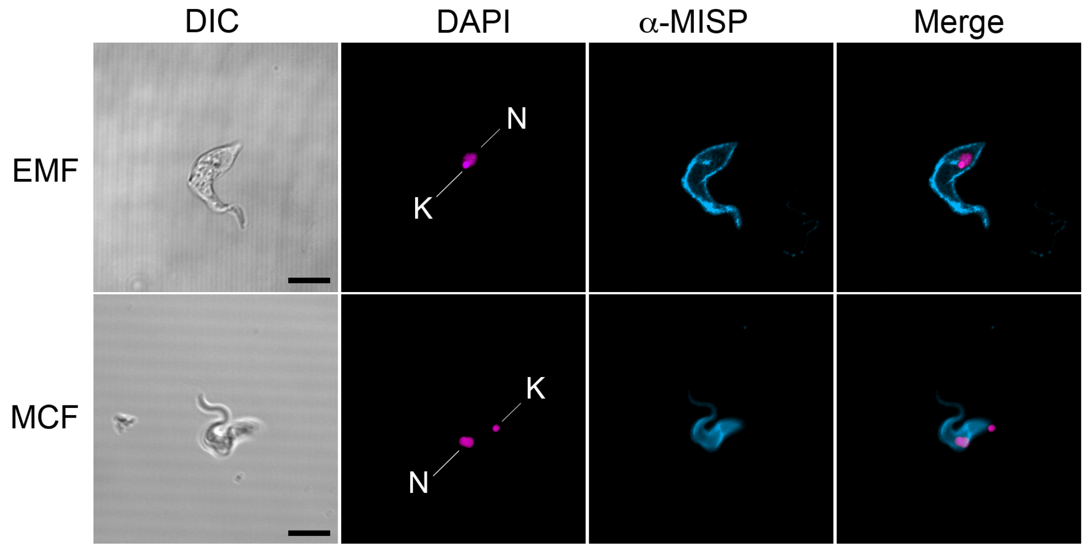

**Supplementary Fig. 8 | Immunodetection of MISP in live SG trypanosomes.** Representative images of live immunostaining of SG parasites using anti-MISP. Epimastigote (EMF) and metacyclic form (MCF) shown by differential interference contrast (DIC) and stained with anti-MISP (red) and DAPI (blue). Nuclei (N) and kinetoplasts (K) noted in the blue channel. Scale bars: 5 μm.

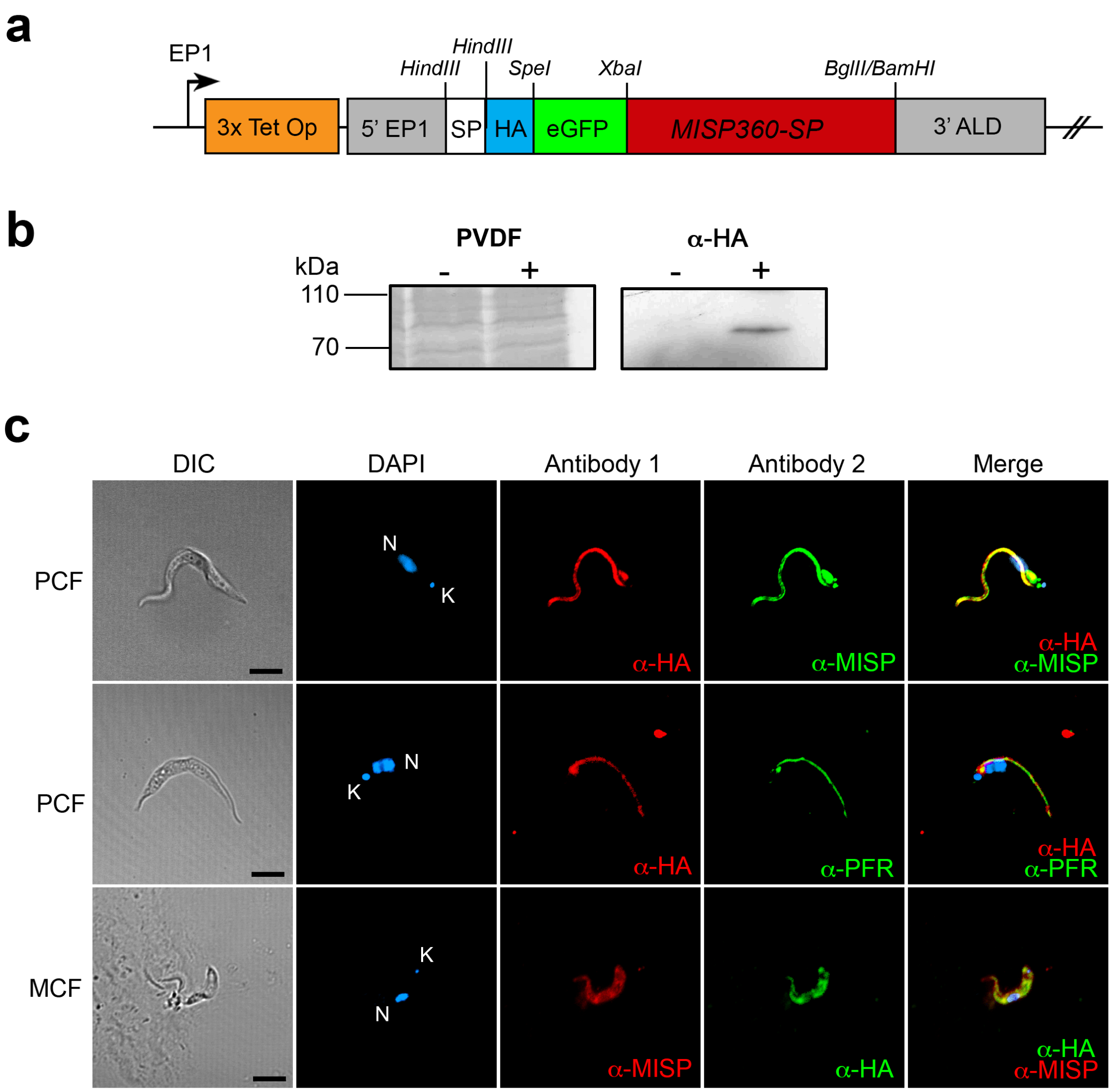

**Supplementary Fig. 9 | Ectopic localization of tagged MISP360. a**, DNA construct encoding for the ectopic tagged *Tb*MISP360. **b,** Immunoblotting for the detection of ectopic (HA-tagged) *Tb*MISP360 expressed by the mutant $^{HA\text{-}eGFP}$MISP360 PCF cell line, either with the transgene uninduced (-) or induced (+), probed with anti-HA (right). PVDF membrane stained with nigrosine after film exposure for sample loading control (left). **c**, Cellular localisation of the HA-tagged ectopic *Tb*MISP360 in AnTat 1.1 90:13 procyclic (PCF) and metacyclic form (MCF) shown by differential interference contrast (DIC); detected by immunostaining on PFA-fixed non-permeabilised cells with either anti-HA plus anti-MISP (co-localisation of ectopic tag with MISP), or with anti-HA plus anti-PFR (flagellar marker) in methanol-fixed cells. All Dox- PCF were negative for both anti-HA and anti-MISP antibodies (not shown). Scale bars: 5 µm.

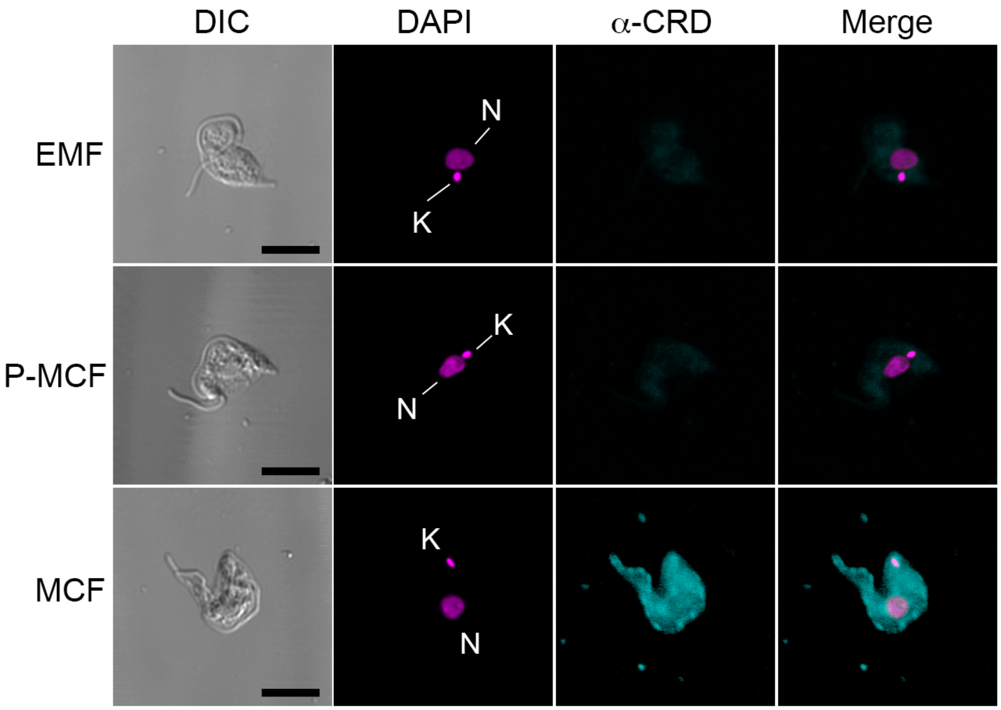

**Supplementary Fig. 10 | Indirect detection of mVSGs in *T. b. brucei* SG stages.** Immunostaining of epimastigotes (EMF), pre-metacyclics (P-MCF) and metacyclics (MCF) from tsetse infected SG with polyclonal anti-CRD antibodies. Nuclei (N) and kinetoplastids (K) noted in the DAPI channel. Differential interference contrast (DIC), DAPI (magenta) and anti-CRD (cyan). Scale bars: 5 µm.

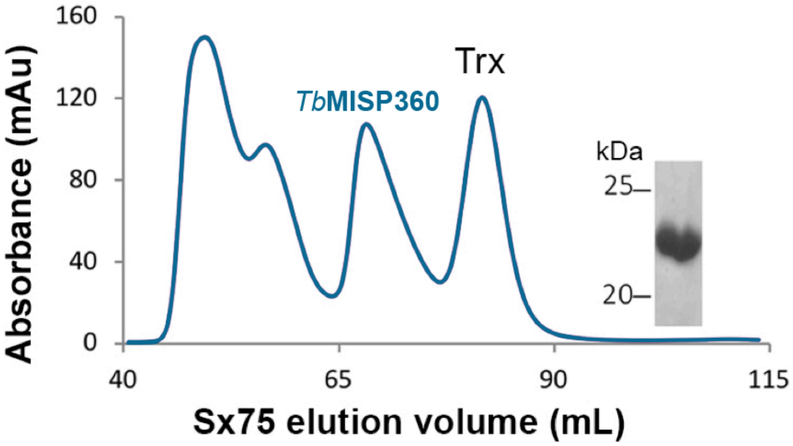

**Supplementary Fig. 11 | SEC elution pattern of recombinant *Tb*MISP360**. Superdex 75 column size exclusion chromatogram of recombinant *Tb*MISP360 and SDS-PAGE analysis of the column fraction (inset) with the protein migrating at ~22 kDa.

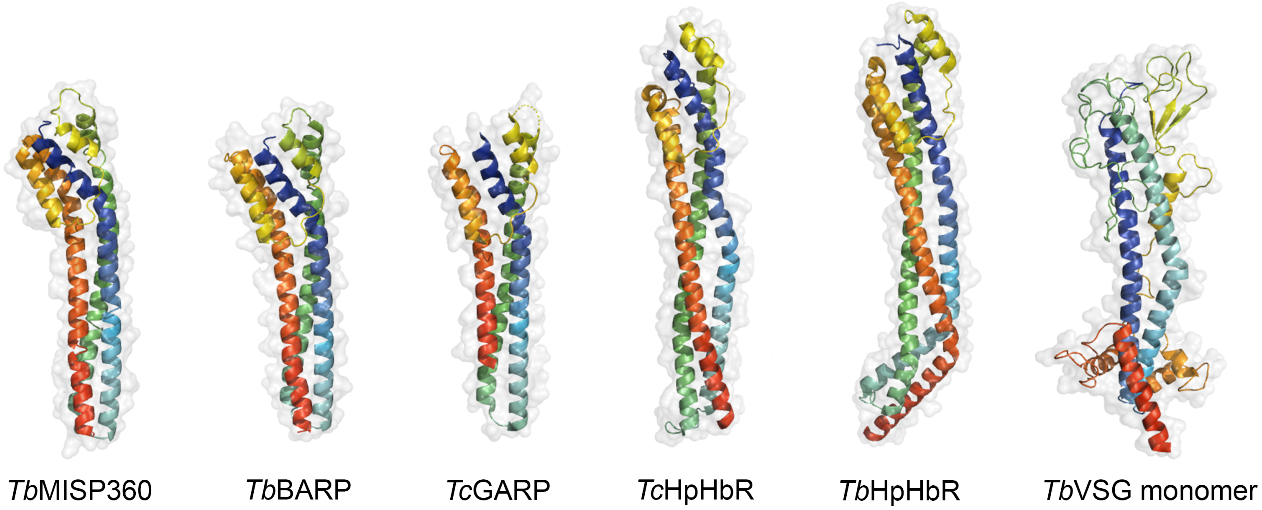

**Supplementary Fig. 12 | Structural analogs of *Tb*MISP360.** Comparison of the crystal structure of the *T. b. brucei* MISP360 N-terminus (*Tb*MISP360) with other trypanosome surface proteins identified as structural analogues; *Tb*MISP360 (PDB: 5VTL), BARP (high confidence model), GARP (PDB: 2Y44), *Tc*HpHbR (PDB: 4E40), *Tb*HpHbR (PDB: 4X0J), and *Tb*VSG 221 monomer (PDB: 1VSG). Structures coloured from blue (N-terminus) to red (C-terminus); molecular surface in semi-transparent grey.

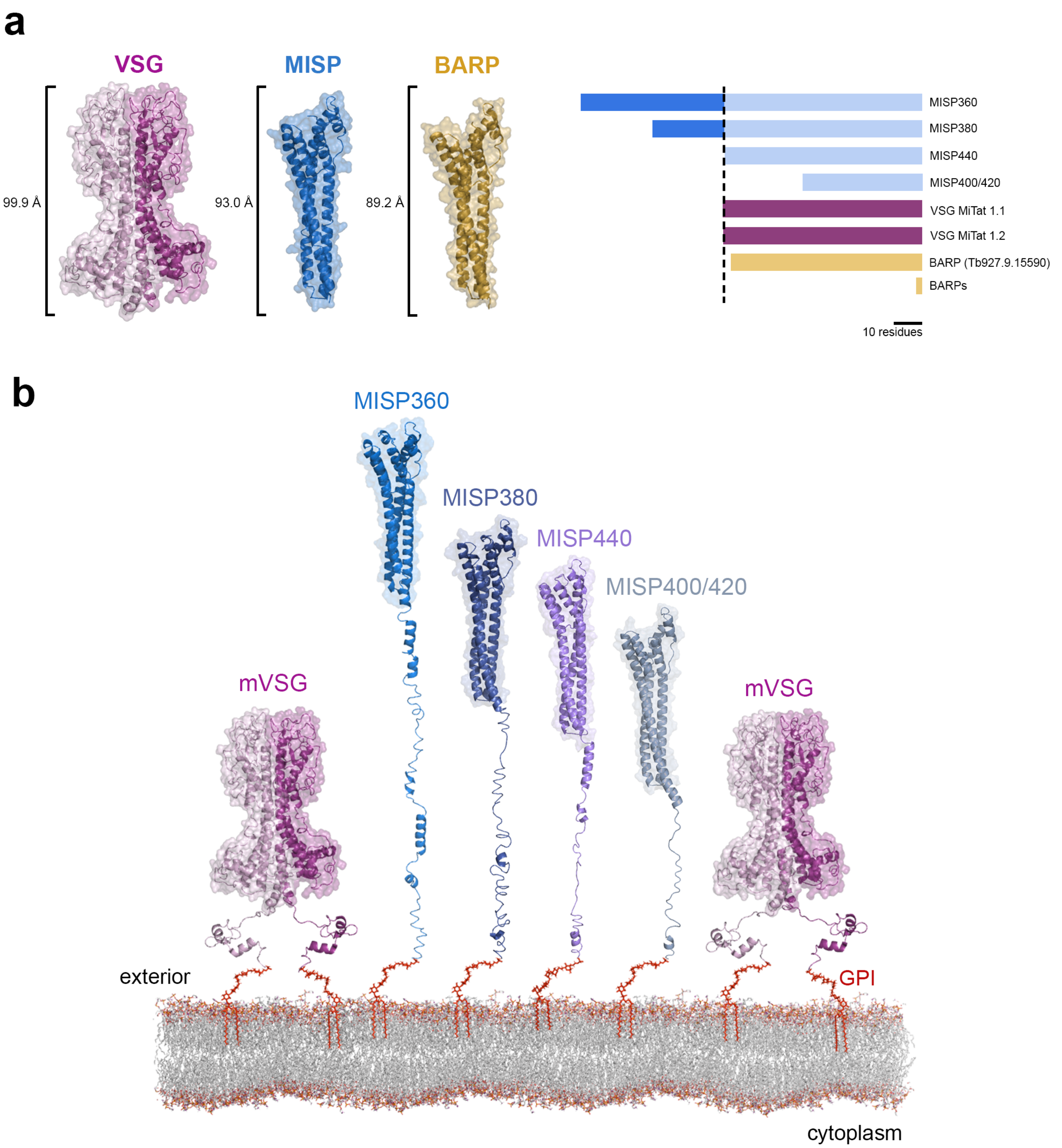

**Supplementary Fig. 13 | Models of MISP C-termini domains. a**, Height comparison between the crystal structure of VSG MiTat 1.2 N-terminus (PDB: 1VSG), the *Tb*MISP360 N-terminus and the BARP model (left). Schematic of the C-terminus domain length of two VSGs, BARP Tb927.9.15590 (the only BARP with C-terminus domain), and the five *T. brucei* MISP (right). **b**, Models of the five *T. brucei* MISP isoforms on the metacyclic surface. mVSGs modelled using the mVAT4 protein sequence and the structures of VSG MiTat 1.2 N- (PDB: 1VSG) and C-terminus (PDB: 1XU6). MISP modelled using the crystal structure of *Tb*MISP360 N-terminus.

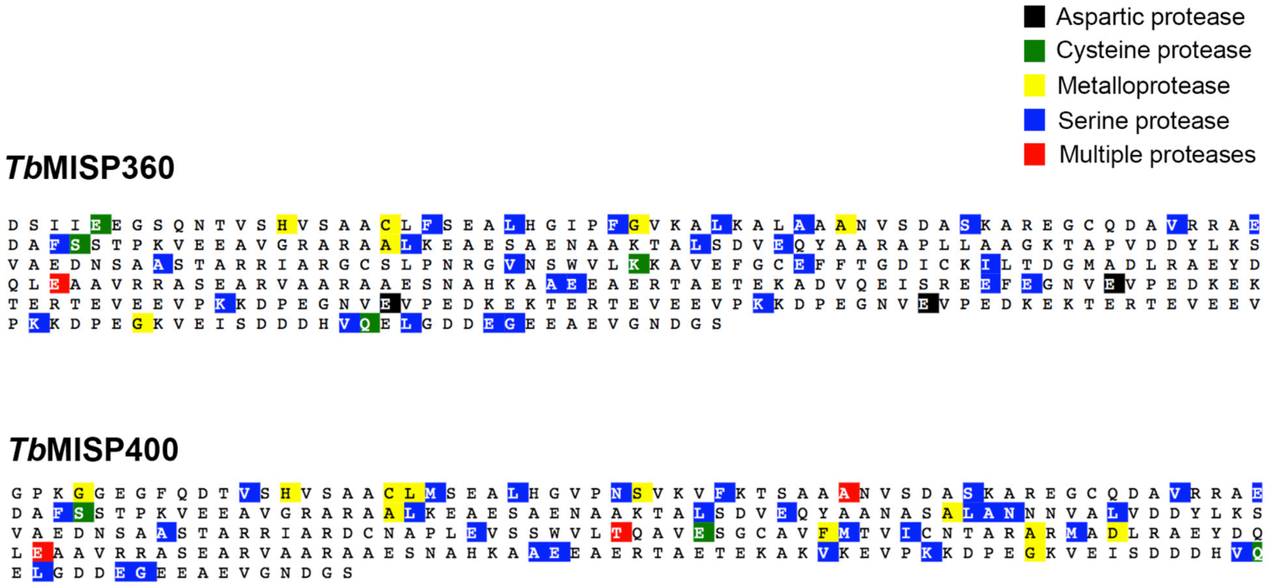

**Supplementary Fig. 14 | PROSPER protease cleavage site predictions.** *Tb*MISP360 (top) and *Tb*MISP400 (bottom) amino acidic sequences with predicted protease cleavage sites highlighted (see key for protease types). Only one member of each MISP subfamily is represented.

**Supplementary Table 1 | Intracellular *T. b. brucei* proteins identified by nLC-MS/MS in infected tsetse saliva.** All identified proteins and peptides have >95% confidence and were manually curated. Accession code (Protein ID), protein description (Annotation), number of identified unique peptides (Peptides), and number of unique spectra (Spectra) are indicated.

| Protein ID [(a)] | Annotation | Peptides | Spectra |
|---|---|---|---|
| Tb927.1.2390 | Beta tubulin | 8 | 12 |
| Tb927.11.11330 | Heat shock protein 70 | 4 | 4 |
| Tb927.10.5620* | Fructose bi-phosphate aldolase | 2 | 2 |
| Tb927.9.6210* | Arginine kinase | 2 | 2 |
| Tb927.10.4560 | Elongation factor 2 | 2 | 2 |
| Tb927.10.10280* | Microtubule-associated protein | 2 | 2 |
| Tb927.10.10460* | Histone 2B, putative | 2 | 2 |
| Tb927.11.3510 | Peptidylpropyl isomerase | 1 | 1 |
| Tb927.3.4850 | Enoyl-CoA hydratase, mitochondrial, putative | 1 | 1 |
| Tb927.11.2470 | Metallo-peptidase, Clan MF, Family M17 | 1 | 1 |
| Tb927.4.2740 | p-25 alpha, putative | 1 | 1 |
| Tb927.10.10890* | Heat shock protein, putative | 1 | 1 |
| Tb927.2.2440* | Proteasome regulatory non-ATPase subunit 6 | 1 | 1 |
| Tb927.10.6510* | Chaperonin HSP60, mitochondrial | 1 | 1 |
| Tb927.4.1340 | Cleavage and polyadenylation specific factor subunit, putative | 1 | 1 |
| Tb927.8.6370 | Cytoskeleton associated protein, putative | 1 | 1 |
| Tb927.3.5530 | Tb-292 membrane associated protein | 1 | 1 |

[(1)]: Accession code of coding gene in strain TREU927, TriTrypDB.
*: Representative ID of multiple homologs sharing the detected peptide.

**Supplementary Table 2 | *In silico* predictions based on MISP amino acid sequences.** Accession code (Protein ID), trypanosome species having the coding gene (Species), MISP sub-family (Sub-family), predicted signal peptide (SP), predicted GPI anchor peptide (GPI), number of predicted *N*-glycosylation sites (*N*-glycan) and number of C-terminus 26 residues motifs (C- mot.) are indicated.

| **Protein ID** [(a)] | **Species** | **Sub-family** | **SP** [(b)] | **GPI** [(c)] | ***N*-gly** | **CTR** [(d)] |
|---|---|---|---|---|---|---|
| Tb927.7.360 | *T. b. brucei / rhodesiense* | MISP-A | 1-17 | Yes | 1 | 3 |
| Tb927.7.380 | *T. b. brucei / rhodesiense* | MISP-A | 1-17 | Yes | 1 | 2 |
| Tb927.7.400 | *T. b. brucei / rhodesiense* | MISP-B | 1-24 | Yes | 1-2 | 0 |
| Tb927.7.420 | *T. b. brucei / rhodesiense* | MISP-B | 1-24 | Yes | 0 | 0 |
| Tb927.7.440 | *T. b. brucei / rhodesiense* | MISP-A | 1-17 | Yes | 1 | 1 |
| Tbg972.7.270 | *T. b. gambiense* | MISP-B | 1-17/24 | Yes | 1-2 | 0 |
| Tbg972.7.290 | *T. b. gambiense* | MISP-A | 1-17/24 | Yes | 1 | 1 |
| TevSTIB805.7.300 | *T. evansi* | MISP-A | 1-17 | Yes | 0 | 3 |
| TevSTIB805.7.320 | *T. evansi* | MISP-A | 1-17 | Yes | 1 | 2 |
| TevSTIB805.7.380 | *T. evansi* | MISP-A | 1-17 | Yes | 1 | 1 |
| TcIL3000.0.02370 | *T. congolense* | - | 1-22 | Yes | 6-7 | - |

[(a)]: Accession code of coding genes in TriTrypDB.
[(b)]: Starting-ending residues. Peptide cut after ending residue.
[(c)]: All GPI anchor peptides are formed by the last 26 residues of the protein.
[(d)]: 26 residues motifs shown in Supplementary Fig. 5a

**Supplementary Table 3 | Summary of *T. brucei* VSG species found in either cultured or fly-derived metacyclics**

| Source [a,b] | RNA (RNA-seq) [c] | Protein (MS/MS) [c] | Reference |
|---|---|---|---|
| *Tbb*-infected tsetse salivary glands (RUMP 503) | Tb927.5.3990 (Q57Z50_TRYB2) | - | Telleria et al. 2014 |
| *Tbb*-infected tsetse salivary glands (RUMP 503) | **VSG ILTat 1.22 (VSI2_TRYBB)** | - | Savage et al. 2016 |
| | **VSG ILTat 1.61 (O97352_9TRYP)** | | |
| | **VSG ILTat 1.63 (Q8MPG1_9TRYP)** | | |
| | **VSG ILTat 1.64 (Q8MPG0_9TRYP)** | | |
| Saliva from *Tbb*-infected tsetse (EATRO 1125 AnTaR 1) | - | **mVAT5 (Q26842_9TRYP)** | Kariithi et al. 2016 |
| | | VSG 1228 (A0A1J0R6Q7_9TRYP) / VSG 1255 (A0A1J0R6U9_9TRYP) | |
| | | VSG 725 (M4SU87_9TRYP) | |
| Cultured *T. brucei* MCF (Lister 427 29:13 *rbp6* overexpressor) | **VSG 397 (A0A1J0R4A4_9TRYP)** | **VSG 397 (A0A1J0R4A4_9TRYP)** | Christiano et al. 2017 |
| | **VSG 531 (M4SYA9_9TRYP)** | **VSG 531 (M4SYA9_9TRYP)** | |
| | **VSG 639 (M4TDP9_9TRYP)** | **VSG 639 (M4TDP9_9TRYP)** | |
| | **VSG 653 (M4SYN2_9TRYP)** | **VSG 653 (M4SYN2_9TRYP)** | |
| | **VSG 1954 (M4T0T6_9TRYP)** | **VSG 1954 (M4T0T6_9TRYP)** | |
| | Tb927.5.4690 / Tb927.9.1050 / Tb927.11.18330 [d] | Tb927.5.291b (B2ZWB3_TRYB2) | |
| | Tb927.1.5300 [e] | Tb927.9.7380 / Tb927.1.5060 [f] | |
| Saliva from *Tbb*-infected tsetse (TSW-196) | - | **mVAT4 (O76421_TRYBR)** | This work |
| | | VSG 221(VSM2_TRYBB) | |
| | | VSG 4959 (A0A1J0RB71_9TRYP) | |
| | | VSG 3088 (M4SWM0_9TRYP) | |
| | | VSG 408/646/769/3613/ 474/1142/4207/4707 (M4SWZ7_9TRYP) | |

**Metacyclic VSGs in bold font**
[a] *In vivo* samples isolated from infected *G. m. morsitans* flies
[b] *T. brucei* strain in parentheses
[c] VSG species; TritrypDB gene accession code; UniProt protein accession code
[d] Genes with higher differential expression (MCF compared to PCF) than canonical mVSGs (>1100 fold)
[e] There are 11 more VSG genes with differential expression MCF/PCF within the canonical mVSG range (250 to 730 fold)
[f] Also upregulated in RNA-seq data

**Supplementary Table 4 | Data collection and refinement statistics of *Tb*MISP360 crystal structure**

| | *Tb*427.07.360 |
|---|---|
| A. Data collection | |
| Synchrotron source | CLS |
| Space group | $P2_12_12_1$ |
| *a*, *b*, *c* (Å) | 24.81, 79.50, 108.17 |
| $\alpha = \beta = \gamma$ (°) | 90.00 |
| Wavelength (Å) | 0.9795 |
| Temperature (K) | 100 |
| Resolution range (Å) | 44.72-1.82 (1.92–1.82) |
| Measured reflections | 136043 |
| Unique reflections | 19605 (2715) |
| Redundancy | 6.9 (6.4) |
| Completeness (%) | 98.2 (94.9) |
| $I/\sigma(I)$ | 20.8 (9.3) |
| $R_{merge}$[a] (%) | 6.0 (15.9) |
| B. Refinement Statistics | |
| Resolution (Å) | 37.31-1.82 (1.88–1.82) |
| $R_{cryst}$[b] / $R_{free}$[c] (%) | 16.94(19.24)/20.28(29.20) |
| No. of atoms | |
| Overall | 3,347 |
| Protein | 3,074 |
| Solvent/Heterogen atoms | 273 |
| Mean temperature factor (Å$^2$) | |
| Overall | 12.3 |
| Protein | 11.3 |
| Solvent/Heterogen atoms | 18.0 |
| r.m.s. deviation from ideality | |
| Bond lengths (Å) | 0.010 |
| Bond angles (°) | 1.14 |
| Ramachandran statistics | |
| Most favored (%) | 99.5 |
| Allowed (%) | 6.6 |
| Generously allowed (%) | 0.0 |
| Disallowed (%) | 0.0 |

Values in parentheses are for the highest resolution shell

[a] $R_{merge} = \sum_{hkl} \sum_i |I_{hkl,i} - [I_{hkl}]| / \sum_{hkl} \sum_i I_{hkl,i}$, where $[I_{hkl}]$ is the is the average of symmetry related observations of a unique reflection

[b] $R_{cryst} = \sum|F_{obs} - F_{calc}| / \sum F_{fobs}$, where $F_{obs}$ and $F_{calc}$ are the observed and the calculated structure factors, respectively.

[c] $R_{free}$ is R using 5% of reflections randomly chosen and omitted from refinement

[d] Ramachandran statistics were determined using PROCHECK
